# Super-Resolution Axial Imaging for Quantifying Piconewton Traction Forces in Live-cells

**DOI:** 10.1101/2024.10.01.615997

**Authors:** Dong-Xia Wang, De-Ming Kong, Jörg Enderlein, Tao Chen

## Abstract

Cell mechanics play a pivotal role in regulating numerous biological processes. While super-resolution microscopy enables the imaging of cellular forces in the lateral dimension with sub-10-nanometer resolution, achieving comparable resolution along the axial dimension remains a significant challenge. In this study, we combine metal-induced energy transfer (MIET) imaging with novel DNA-hairpin-based molecular tension probes (MIET-MTP) to map integrin-mediated mechanical forces with nanometer precision in the axial direction. MIET-MTP allows for the simultaneous observation of both the plasma membrane and force-exerting molecules in the axial dimension. Using this approach, we mapped axial integrin tension in focal adhesions and podosomes, alongside their corresponding plasma membrane height profiles, offering detailed insights into the structures involved in force transmission.

---

Cell mechanics play a crucial role in regulating various biological processes, such as immune recognition, blood coagulation, cell migration, and differentiation [1–3]. Structures responsible for force transmission, including filopodia, focal adhesions, podosomes, and the cytoskeleton, are dynamically organized at the nanoscale in all dimensions [4, 5]. To gain a deeper understanding of how mechanical forces interact with biochemical signaling pathways, it is essential to develop advanced methods capable of mapping the distribution of forces within living cells. Molecular tension probes (MTPs) have been developed to measure receptor forces with piconewton (pN) sensitivity [6, 7]. These probes consist of an extendable “spring-like” element, such as polyethylene glycol or DNA hairpins, flanked by a fluorophore and a quencher, and with one end immobilized on a surface. When receptor forces are applied to the other end of the probe, the “spring” extends, causing the fluorophore and quencher to separate, which in turn increases fluorescence. This change in fluorescence can be detected using various microscopy techniques, enabling the mapping of receptor forces in cellular adhesion structures such as podosomes [8, 9], focal adhesions (FAs) [10, 11], or E-cadherin complexes [12].

Current fluorescence techniques, such as total internal reflection fluorescence microscopy, confocal microscopy, and structured illumination microscopy, are limited to mapping forces in the lateral dimension [13, 14]. Even with super-resolution techniques like localized super-resolution microscopy like DNA points accumulation for imaging in nanoscale topography (DNA-PAINT), which can achieve resolutions of a few nanometer in the lateral direction [15, 16], the resolution in the axial dimension is limited to tens of nanometers [15]. However, because cells apply axial forces to the MTPs to separate the fluorophore-quencher pair, accurately characterizing tension forces along the axial direction is more critical than in the lateral plane. Despite this need, no current technique allows for mapping cellular forces with nanometer resolution in the axial direction.

MIET imaging uses the phenomenon that a fluorescence emitter, when brought close to a metal surface, transfers its excited-state energy to surface plasmons (collective oscillations of free electrons) in the metal [17–19]. This energy transfer is distance-dependent, resulting in the modulation of both fluorescence lifetime and intensity based on the proximity between the emitter and the metal surface. This mechanism is analogous to Förster resonance energy transfer (FRET), which involves energy transfer from a donor to an acceptor fluorophore. The measured fluorescence lifetime can then be converted into a precise distance between the emitter and the metal surface using semiclassical quantum-electrodynamic theory [20] (see Methods). Additionally, due to the broad absorption spectra of metal, the energy transfer from a fluorescent molecule to the metal film occurs with high efficiency across the visible spectrum, allowing any dye within this spectral range to exhibit the effect. This feature facilitates the co-localization of multiple fluorophores through MIET (Figure 1a).

**Fig 1.**
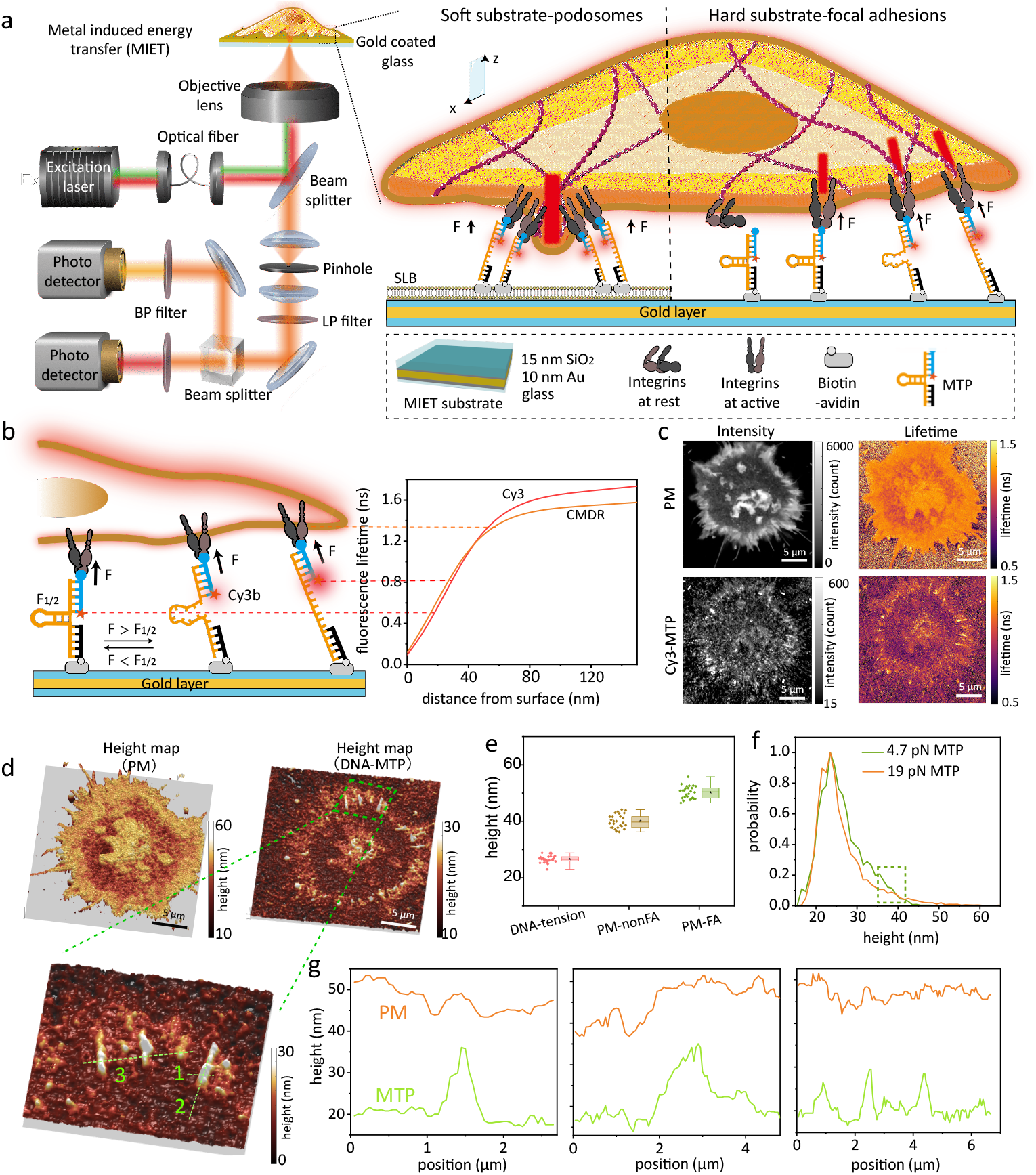
MIET-MTP measurement on FA. (a) Left: diagram showing the MIET-MTP setup for two-color measurement. Cos7/NIH 3T3 cells are seeded on the MTP-modified MIET substrate, which consists of a 10-nm gold film sandwiched between a 15-nm silica layer and a glass coverslip; Right: schematic illustrating integrin-involved adhesion structures: podosome and FA. (b) Working principle of MIET-MTP, which includes an anchor strand immobilized on the MIET substrate, a hairpin strand that unfolds under sufficient tension and a ligand stand presenting an adhesive peptide. The right panel shows the calculated fluorescence lifetime-versus-distance curves (MIET curves) for fluorophores Cy3 and CMDR fluorophores, respectively. The calculation details provided in Supplementary Table 2. (c) The fluorescence intensity images and corresponding lifetime images used to determine the heights of Cy3 and CMDR, respectively. (d) Reconstructed height maps for MTP and PM. The bottom panel shows the enlarged area. (e) Box plot of the height values for MTP, PM without FA (PM-nonFA), and PM with FA (PM-FA). Box plots show the 25th–75th quantiles (box), median (solid line), mean (black dot), and whiskers (minima to maxima). n = 26 independent cells. (f) Histogram of MTP heights for 4.7 pN and 19 pN MTP, respectively. (g) Linescan analysis for the height profiles of PM and MTP.

Matrix-activated integrins can form different adhesion structures depending on the substrate [21]: focal adhesions (FAs) when spread on rigid surfaces, and podosomelike adhesions on fluid lipid surfaces (Figure 1b). We employed MIET-MTP to study both types of adhesion structures. To map forces along the axial direction, we utilized MIET in conjunction with a widely recognized DNA-based hairpin MTP (Figure 1b).

One end of the MTP was modified with an Arg-Gly-Asp peptide (RGD), designed to activate and bind integrin on the cell surface. The other end was modified with a biotin molecule to anchor the MTP to an avidin-modified surface or an avidin-modified supported lipid bilayer (SLB) on a MIET substrate. The MIET substrate comprises a 10-nm gold film sandwiched between a 15-nm silica layer and a glass coverslip (Figure 1a). Since the gold film acts as a quencher in MTPs, no additional quencher molecule is required.

When integrin-mediated forces exceed the *F*_1*/*2_, the equilibrium force at which 50% of hairpins unfold, the dye lifts from the surface, leading to an increase in fluorescence lifetime and intensity (Figure 1b). In contrast, hairpins experiencing forces lower than *F*_1*/*2_ remain folded and positioned closer to the surface, exhibiting a shorter fluorescence lifetime and intensity. We followed previous reports using DNA-MTPs with *F*_1*/*2_ values of 4.7 pN or 19 pN [11] (Supplementary Table 1), both of which are below the reported force threshold for initial integrin adhesion [22], estimated to be ∼ 40 pN. Additionally, by labeling the plasma membrane (PM) with a separate fluorophore, we reconstructed the height profile of the basal PM decorated with MTPs (Figure 1b), providing insights into how adhesion structures affect the basal membrane.

Initially, we tested our ability to measure the traction forces generated by focal adhesions in Cos7 cells, as mechanical forces regulate their activation and clotting [5]. Cos7 cells were seeded on Cy3-MTPs-modified MIET substrate, which produced a robust signal corresponding to integrins applying forces greater than 4.7 pN (*F*_1*/*2_). The PM was labeled with a commercial dye, CellMask Deep Red (CMDR), to report its height. Control experiments confirmed that no FRET occurs between Cy3-MTPs and CMDR due to the significant distance between the Cy3 position and the PM (Supplementary Note 2, Supplementary Figure 5). This setup anabled us to simultaneously measure Cy3-MTPs and PM height using two detectors (Figure 1a).

We imaged the sample using a laser-scnning confocal microscope (CLSM) equipped with time-correlated single-photon counting (TCSPC) to perform fluorescence lifetime measurements. To accurately determine the lifetime values for both the fluorophores in the MTPs and the PM, we scanned individual cells to accumulate sufficient signals for reliable fluorescence decay fitting. TCSPC curves were generated for each pixel and fitted with a multiexponential decay model (Supplementary Note 3), yielding a mean fluorescence lifetime for each pixel (50 nm *×* 50 nm). These lifetime images were subsequently converted to height maps above the silica surface using fluorophore-specific MIET calibration curves.

Figure 1c shows the measured fluorescence intensity and lifetime images for MTPs and PM in a single Cos7 cell after 20 min of incubation. The cell edges exhibit higher intensity (approximately 10 times stronger than the background) and longer lifetime for both MTPs and PM, indicating a higher density of hairpin unfolding at these regions. This observation is consistent with previous reports using MTPs based on fluorophore-quencher pairs [11, 23]. We further constructed 3D height maps for MTPs and PM (Figure 1d) from the lifetime images. FA areas are clearly identifiable by their elevated MTP height. We calculated the average height for each FA region, focusing on areas with a signal-to-noise ratio (SNR) greater than 10 (see Supplementary Figure 6). The mean height of the FAs is approximately 26 nm, which is ∼ 9 nm higher than the height of closed MTPs (17.6 nm, Supplementary Figure 7, Supplementary Note 4). In MIET measurements, the axial localization precision is determined by the number of photons used in the fluorescence lifetime fitting. For DNA-MTPs, the minimum photon count per pixel is approximately 400 (Figure 1c, Supplementary Figure 8, Supplementary Figure 9). Fitting with 400 photons results in a height uncertainty of around 2 nm (Supplementary Note 3). For the PM height, the minimum photon count per pixel is approximately 1500, which produces a height uncertainty of less than 1 nm. It is important to note that the height values obtained for each pixel is a spatial average over the the excitation focus area, encompassing both folded and unfolded MTPs. However, the signal is dominated by the high fluorescence intensity of the unfolded MTPs (unfolded MTPs exhibit at least ten times higher brightness than folded MTPs, Supplementary Figure 13), so that the measured lietime mostly reports about the height of these unfolded MTPs.

We further performed a linescan analysis on both the FA and PM within the same area (Figure 1d and 1g). Interestingly, a positive correlation was observed: higher FAs correspond to higher PM heights. This same trend was also seen in the 19 pN MTPs (Supplementary Figure 10). This positive correlation suggests that integrin-mediated forces unfolds the DNA, simultaneously elevating the membrane. Figure 1e presents the statistical data from 26 cells using 4.7 pN MTPs. Our results demonstrated distinct structures for the FA and PM: the unfolded MTPs have a mean height of 26.6 *±* 1.3 nm (mean *±* SD, N = 26), while the FA regions of the PM had a mean height of 50.2 *±* 2.3 nm, and the non-FA regions of the PM showed a mean height of 40.0 *±* 2.4 nm.

From the MTP height images, we observed that the heights of the unfolded 4.7 pN MTPs varied wildly, ranging from 20 nm to 40 nm (Figure 1f), indicating heterogeneity in the tension forces. Theoretically, the maximum height for fully stretched 4.7 pN MTPs should be approximately 35 nm if the DNA chain is fully extended and remains upright on the surface. However, about 5% of the FA regions displayed heights greater than 35 nm, which could be attributed to DNA overstretching under constant tension [24]. To further investigate this hypothesis, we performed anisotropy measurements on Cy3-MTPs on MIET substrates, which report the rotational motion of the dye molecule during its excited-state lifetime [25]. In anisotropy measurements, polarized light is used to excite the fluorophore, and the emitted light is separated by polarization using a polarization beam splitter (Figure 2a). The polarized emission is then directed onto two detectors, allowing for the calculation of fluorescence anisotropy, *r*, from the intensities detected by these two detectors (Supplementary Figure 11). Cy3 dye can stack against the nucleobases of the DNA duplex, perpendicular to its long axis, which is essential for maintaining a fixed dye orientation relative to the biomolecule, enabling accurate determination of its orientation [26, 27].

**Fig 2.**
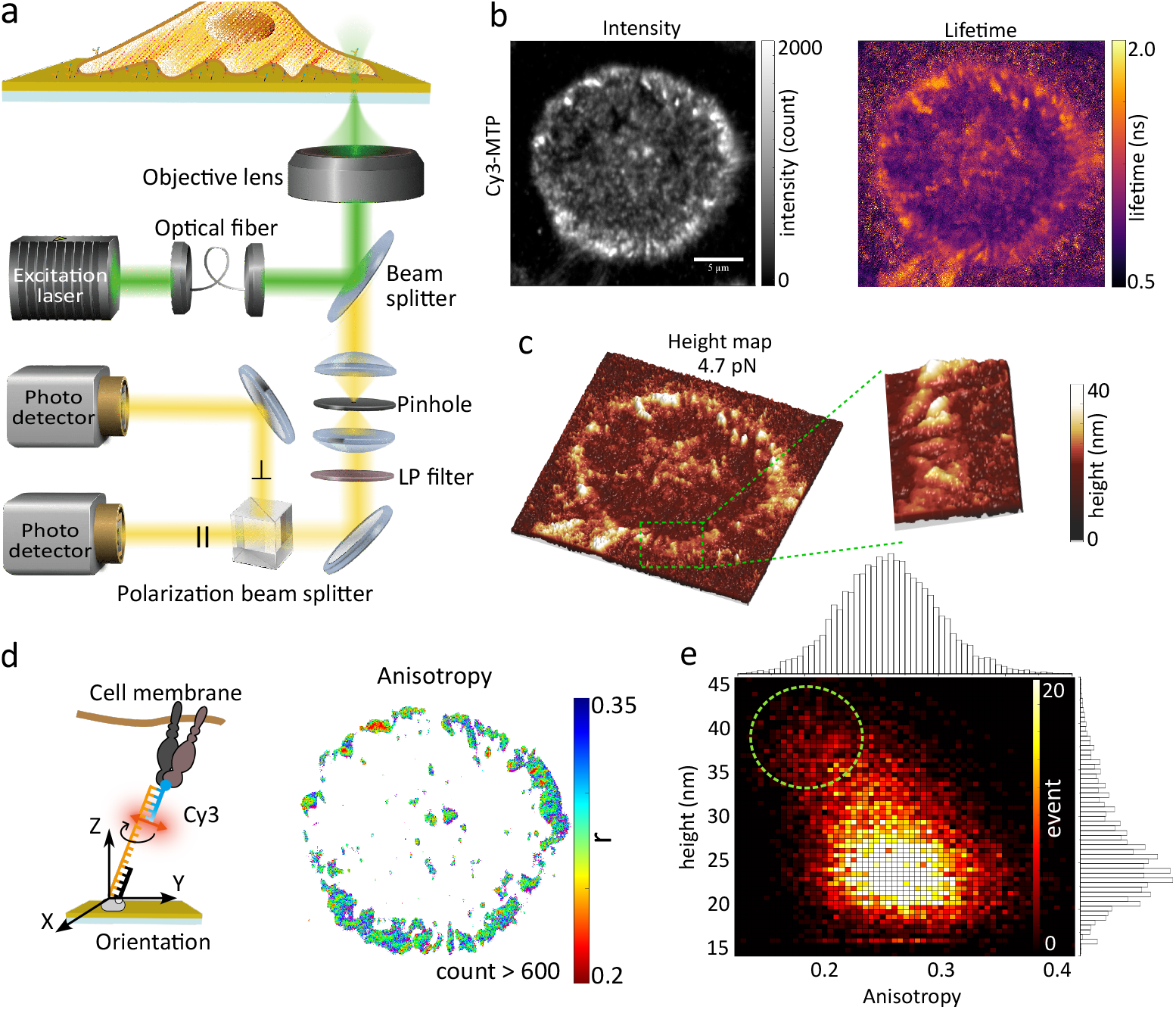
MIET-anisotropy measurement on FA. (a) Diagram illustrating the MIET-anisotropy measurement. The Cy3 on the MTP is excited by polarized light and the emission light is separated based on its polarization using a polarization beam splitter into two detectors. (b) Fluorescence intensity and lifetime image of a cell on 4.7 pN MTP-modifed MIET substrate. (c) The corresponding height map of the MTP. (d) Left: Illustration showing the Cy3 orientation perpendicular to the long axis of the tension probe. Right: Calculated anisotropy image of Cy3 on MTP. (e) 2D histogram depicting the relationship between the height and anisotropy.

Figure 2 shows the anisotropy measurements for a single cell on MTP-MIET substrate. The formation of FAs is evident from the intensity, lifetime, and height images (Figure 2b and 2c). As expected, we observed high anisotropy (mean *r* ∼ 0.27 for the FAs) in the anisotropy image (Figure 2d), indicating that the rotation of Cy3 is restricted due to stacking on the DNA duplex. Interestingly, a bias towards lower anisotropy values was found at greater heights in the 2D height-anisotropy histogram (Figure 2e). The lower anisotropy suggests increased flexibility of the fluorophore, indicating a reduction in the stacking of Cy3 on the DNA duplex at higher elevations. This increased flexibility is likely caused by DNA overwinding when stretched. Additionally, no bias effect was observed for probes using 19 pN Cy3-MTPs (Supplementary Figure 12), as the 19 pN MTP has a longer DNA stretch length in the unfolded state (∼ 45 nm). These MIET-anisotropy measurements provide further insight into the force structures of FAs using MTPs, suggesting that caution is necessary when using DNA for orientation measurements, particularly under varying force conditions.

Next, we mapped podosome-like adhesions and the PM on supported lipid bilayers (SLBs) using MIET-MTP (Figure 3a). Previous studies have shown that integrins can apply ring-like tensile forces in podosomes on SLB [8, 9] (Figure 3b). Consistent with these findings, we observed ring-like structures in the MTP fluorescence intensity images, and the corresponding podosomes exhibited lower heights in the PM height maps (Figure 3c). We found that regions with longer lifetimes in the MTP lifetime image correlated with higher intensities in the MTP intensity image. However, the ring-like structures were difficult to discern in the MTP lifetime or height images (Figure 3c and 3d), likely due to the low SNR. In the podosome measurements, the SNR was approximately 2, which led to an underestimation of lifetime values (Supplementary Note 5, Supplementary Figure 13). The low SNR may result from factors such as fluorophore diffusion, minimal height increases, and a low percentage of open MTPs. To confirm that the observed increase in lifetime is due to MTP unfolding, we conducted a control experiment in which cells were seeded on SLBs with MTPs that lacked RGD ligands (Supplementary Figure 14). Interestingly, even without the RGD, increased intensity was observed in the MTP intensity image, but no corresponding increase in lifetime was detected, indicating that the MTPs remained folded. Nevertheless, podosome formation was confirmed through PM measurements. The increased intensities in the MTP image likely resulted from MTP clustering around the podosomes, as the MTPs are free to diffuse within the SLB (Supplementary Figure 15).

**Fig 3.**
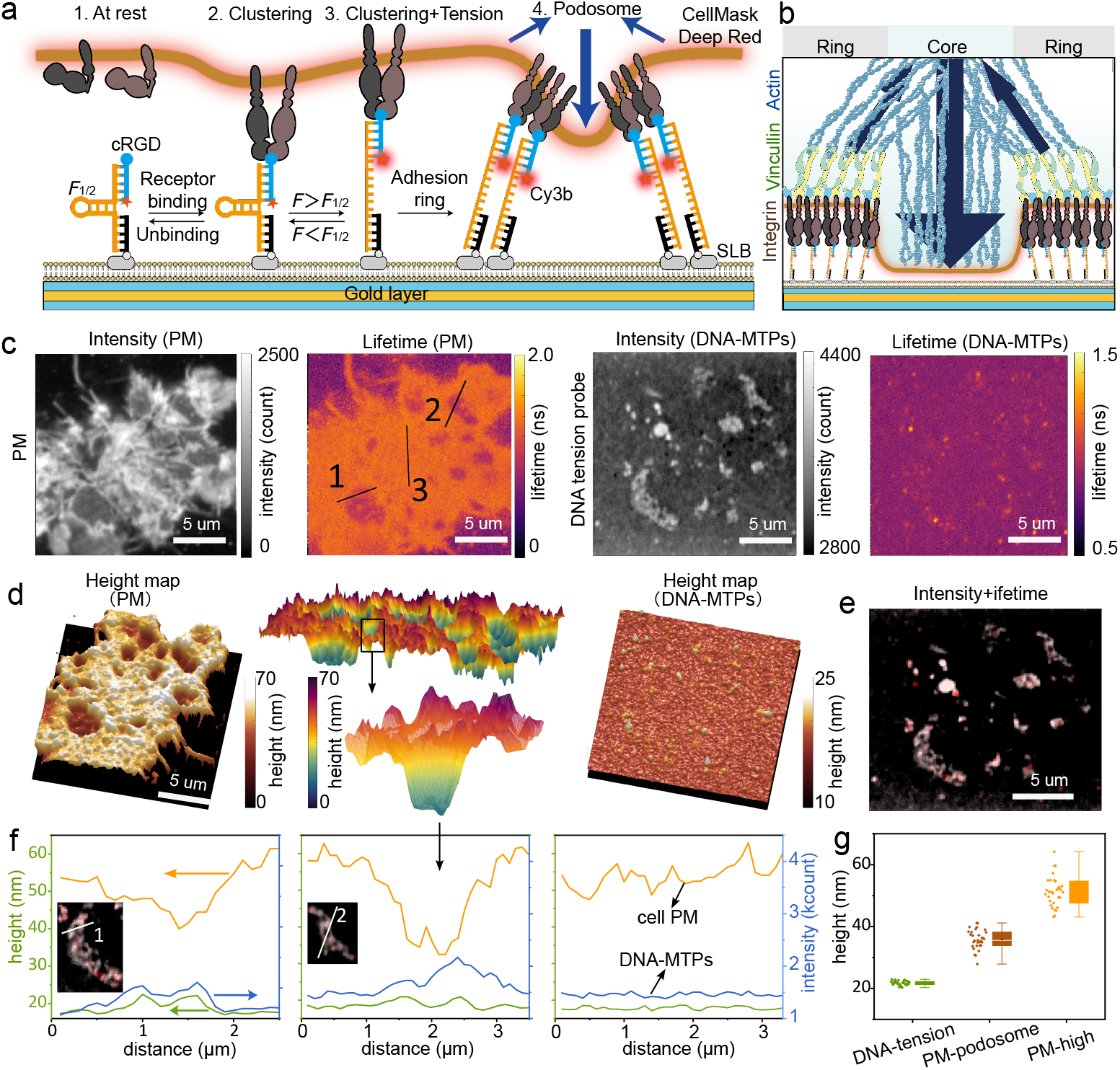
MIET-MTP measurement on podosomes. (a) Diagram illustrating the formation of podosomes in NIH 3T3 fibroblasts. The cells are seeded on the MTP-modified SLB, which is supported on a MIET substrate. (b) Schematic of a single podosome at the cell–SLB interface. (c) Fluorescence intensity images and corresponding lifetime images for the heights of Cy3 and CMDR, respectively. (d) The reconstructed height maps for MTP and PM. The PM heihgt maps are shown in top view (left) and bottom view (middle). (e) Overlay of the MTP intensity image with MTP lifetime image. (f) Linescan analysis for the lines marked in **c**. (g) Box plot of the height values for MTP, PM-podosome, and PM without podosome (PM-high). n = 30 independent cells.

Importantly, the podosome structures were clearly visible in the PM height profiles (Figure 3d). A negative correlation between the unfolding MTPs and PM was observed in the podosome measurements, as shown by the linescan analysis (Figure 3f). Podosomes exhibited lower PM height, which consistently corresponded to higher DNA probe heights and greater brightness. Statistical analysis of 30 cells revealed that the unfolding MTPs had a mean height of 21.7 *±* 0.7 nm, the podosome-associated PM (podosome-PM) had a mean height of 35.8 *±* 3.1 nm, and the non-podosome PM areas had a mean height of 51.3 *±* 4.9 nm (Figure 3g). Control experiments showed that the folded MTPs on SLBs had a mean height of 17.1 *±* 1 nm from the surface (Supplementary Figure 7), indicating only a 4.6 nm increase in height after MTP unfolding.

It is important to note, due to the low SNR, the actual height of MTPs is likely higher than the measured values. The negative correlation observed is explained by the fact that the core of the podosome structure, where the podosome-PM height is measured, lacks integrins [8, 28] (Figure 3b); instead, the forces are generated by integrins distributed around the ring of the podosome. To determine the PM height at the podosome rim, we calculated the PM height for areas displaying only longer lifetimes in the MTP lifetime image. The mean height for this area (ring-podosome PM) is 44.0 *±* 7.2 nm, higher than the core-podosome PM, though still lower than the non-podosome PM. Measurements using 19 pN MTPs also showed the same negative correlation between the unfolded MTPs and podosome-PM, suggesting that forces generated within podosomes exceed 19 pN (Supplementary Figure 16).

## Discussion

Current techniques for mapping cellular force are primarily focused on the lateral plane. MIET-MTP offers a complementary approach, enabling the resolution of cellular forces along the axial direction with nanometer precision. A key advantage of MIET-MTP is its ability to simultaneously observe the PM and force-exerting molecules, offering detailed insights into the structures involved in force transmission. To demonstrate the capabilities of MIET-MTP, we mapped integrin tension in FA and podosome, revealing structural differences. Notably, we observed distinct behaviors despite the involvement of the same force-bearing molecules: FAs exhibited a positive height correlation between the MTP and PM, while podosomes showed a negative correlation. This difference reflects the unique structures and functions of these adhesion complexes. Additionally, by combining MIET-MTP with anisotropy measurements, we demonstrated that DNA within the MTP molecule becomes overstretched under integrin force.

One limitation of MIET-MTP is its current reliance on ensemble measurements (Supplementary Note 5), which can lead to an underestimation of unfolded MTPs’ height, particularly in cases with low SNR. However, this limitation could be mitigated by integrating DNA-PAINT, which would enable the resolution of single force events. The ability to map forces and associated structures, such as the cytoskeleton, along the axial direction simultaneously is a powerful capability. We anticipate that MIET-MTP will become a standard technique for bridging structural biology with mechanobiology.

## Methods

### Ethical Statement

The research conducted in this study complies with all relevant ethical regulations.

### MTP sensor synthesis and purification

All oligonucleotides (listed in Supplementary Table 1) were ordered from Thermo Fisher, except for the ligand (DBCO-modified ssDNA FAM/Cy3), which was purchased from Biomers.net. A total of 5 nmol of DBCO-ligand and 30 nmol of Azide-cRGD (azide-modified cyclo [Arg-Gly-Asp-D-Phe-Lys(PEG-PEG)]) (Peptides International, cat. no. RGD-3759-PI) were conjugated in 1× PBS (pH 7.4) overnight at 15 ^°^C, using a metal bath at 300 rpm in a total volume of 1 mL. The product was purified using C18 reversed phase column (Thermo Fisher), with a gradient from buffer A (0.1 M TEAA triethylammonium acetate buffer) to buffer B (acetonitrile) over the course of 60 minutes (Supplementary Figure 1). Peak fractions were collected and filtered through ultrafiltration centrifuge tubes with a molecular weight cutoff of 3.5 kDa. Successfully conjugated DNA ligands were stored at -20 ^°^C until further use.

To synthesize MTP probes, three oligonucleotides at equal molar concentrations were mixed in 1× PBS (pH 7.4). The mixtures were annealed by heating to 95 ^°^C, then cooled to 25 ^°^C at a rate of 1 ^°^C/min in a 0.2 mL thermowell tube. To chemically open MTPs, MTPs were hybridized with a 5× molar excess of complementary sequence.

### Small unilamellar vesicle (SUV) preparation

To prepare SUVs, 100 µL of a 10 mg/mL solution of 1,2-dioleoyl-sn-glycero-3-phosphocholine (DOPC) lipids (Sigma-Aldrich, P6354) and 0.05–0.2 mol% biotinylated lipids (1,2-dioleoyl-sn-glycero-3-phosphoethanolamine-N-(biotinyl) (Biotinyl-Cap PE)) (Sigma-Aldrich, 870273P) in chloroform was vacuum-driedg at 30 ^°^C for 1 hour to remove the solvent. Subsequently, 500 µL of PBS buffer (pH 7.4) was added, and the mixture was incubated at 30 ^°^C with shaking for 1 hour. The resulting lipid suspension was then extruded through a 50 nm polycarbonate filter (Whatman) for 15 cycles. These vesicle solutions should be used within 3 days and stored at 4 ^°^C until use.

### MIET and GIET measurements

All measurements were conducted using a self-constructed confocal microscopy. A white laser with Acousto-Optic Tunalble Filters (AOTFnC-400.650-TN, Pegasus Optik GmbH) operating at a repetition rate of 80 MHz served as the excitation source. The laser beam was filtered with a clear-up filter (LF405/488/561/635, Semrock) and collimated through an infinity-corrected 4x objective (UPISApo 4X, Olympus). The polarized beam passed through a quarter wave plate (AHWP10M-600, Thorlab) and was then reflected by a dichroic mirror (Di01-R405/488/561/635, Semrock) towards a high numerical aperture objective (UApoN 100X, oil, 1.49 N.A., Olympus). Fluorescence emission was focused into a pinhole with a diameter of 100 µm and refocused onto two avalanche photodiodes (*τ* -SPAD, PicoQuant). A band-pass filter (Brightline HC692/40, or FF01-609/54-25, or FF01-525/45-25, Semrock) was positioned before each detector, corresponding to the different fluorophores. Single-count data from the detectors were processed using a multi-channel picosecond event timer (Hydraharp 400, PicoQuant). Imaging scanning was facilitated by a fast Galvo scanner (FLIMbee, Picoquant). For live cell measurements, a microscopy-based incubator (ibidi Stage Top Incubation System-Blue Line, ibidi) maintained controlled conditions (temperature: 37 °C, CO_2_ concentration: 5%, humidity: 40%). For anisotropy measurements, a broadband polarizing beamsplitter cube (Thorlab) was placed before the two detectors and the excitation laser was polarized, with its direction aligned parallel to one detector.

### MIET calibration curves calculation

The geometry of a MIET experiment is illustrated in Supplementary Figure 2. A fluorescent molecule is positioned at a distance *z* above a substrate, which consists of a 10-nm-thick gold layer and a 15-nm silica layer on a commercial glass coverslip. Fluorescence excitation and detection are conducted through this substrate from below. For calculating the MIET calibration curve (lifetime versus distance curve), the emitting molecule is treated as an ideal oscillating electric dipole, and its emitted electromagnetic field is mathematically described as a superposition of plane waves. The interaction of each plane wave with the planar substrate is calculated using standard Fresnel theory, yielding the complete electromagnetic field of the emitter in the presence of the substrate. By integrating the Poynting vector two parallel planes enclosing the emitter, the total energy flux of the emitted field can be determined, enabling the calculation of the full emission rate of the dipole. By applying the same approach to two planes enclosing the MIET substrate (the metal layer), the fraction of energy absorbed by the substrate can be calculated. From these calculations, the emission rate *S*(*θ, z*) of an ideal electric dipole is found to be:

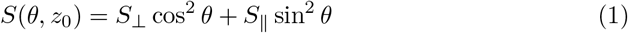

where *θ* represents the angle between the dipole’s axis and the vertical direction (normal to the surface), and the functions *S*_⊥_ and *S*_∥_ depend solely on the orientation *θ*.

Additionally, real fluorophores exhibit nonradiative transitions from the excited to the ground state, which determines the quantum yield *φ* of the fluorophore:

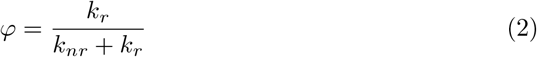

Here, *k*_*r*_ is the radiative transition rate, and *k*_*nr*_ is the nonradiative transition rate. When comparing the fluorescence lifetime of a free molecule far from the substrate to that of a molecule at a distance *z* above the substrate, we get:

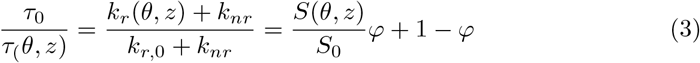

where *S*_0_ represents the free-space emission power of an ideal electric dipole emitter and is given by 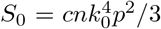, with *c* being the speed of light, *k*_0_ the wavevector in vacuum, *n* the refractive index of water, and *p* the amplitude of the dipole moment vector.

Finally, the fluorescence lifetime *τ*_(*θ,z*)_ as a function of distance *z* is fitted using Equation 3, with the angle *θ*, the free-space lifetime *τ*_0_, and the quantum yield value *φ* as fitting parameters. The free-space lifetimes for all fluorophores were measured using samples on glass surfaces (Supplementary Figure 3). Additional details regarding the determination of fluorescence lifetimes, quantum yields, and fluorophore orientations are provided in Supplementary Note 1. All fitting parameters are listed in Supplementary Table 2, and the calculated MIET curves are shown in Supplementary Figure 4.

### Substrate preparation

The protocol for preparing the gold-modified substrate has been previously described in our publications [17, 29]. Briefly, a layer-by-layer electron-beam evaporation process was used to deposit a 2 nm titanium layer, followed by a 10 nm gold layer, another 1 nm titanium layer, and finally a 15 nm SiO_2_ layer on the surface of a glass coverslip. The deposition rate was kept at the slowest rate (1 °As^-1^) to ensure maximal homogeneity. The spacer thickness was continuously monitored during evaporation using an oscillating quartz unit. This gold-covered substrate is referred to as the MIET substrate.

To prepare the SLB on MIET substrate, the MIET coverslip was first activated for 30 seconds with low-intensity plasma from a plasma cleaner (Harrick Plasma, New York, USA). After activation, a droplet of the SUV solution was placed onto the substrate and incubated for 1 hour to promote the formation of a continuous bilayer with minimal imperfections. The substrate was then thoroughly washed with 1× PBS buffer. Next, the substrate was incubated with Neutravidin (Thermo Fisher Scientific, cat. no. 31000) at a final concentration of 0.5 mg/mL for 20 minutes. After another wash with 1× PBS, the substrate was incubated for 60 minutes with 200 nM of biotin-labeled MTP and subsequently washed again with 1× PBS.

For the hard substrate, after the MIET coverslip was activated, biotin-labeled BSA (Sigma-Aldrich, A8549) at 500 µg/mL concentration was added and incubated for 20 min. After washing with water, neutravidin solution was added directly without the addition of SUVs. The remaining steps were identical to those used for preparing the soft substrate.

### Cell culture and imaging

Cells were cultured in Dulbecco’s Modified Eagle Medium (DMEM) supplemented with 10% (v/v) fetal bovine serum and 1% antibiotics (penicillin-streptomycin-amphotericin B) until they reached approximately 75% confluency. The cells were then detached from culture flasks using a 0.25% (wt/vol) trypsin solution (Corning). The prepared cells were seeded onto MTP-functionalized surfaces at a density of 20,000 cells/cm^2^. Imaging of the cells was conducted between 20 and 120 minutes post-plating.

### Statistics & Reproducibility

All values are expressed as the mean *±* SD. Box plots show the 25th–75th quantiles (box), median (solid line), mean (black dot), and whiskers (minima to maxima). No statistical method was used to predetermine sample size. All experiments were repeated at least forth with reproducible results.

## Data Availability

Source data are provided with this paper. There are no restrictions on data availability. All data supporting the findings of this study are available within the main text, supplementary information, and ‘Source data’ Excel file.

## Code Availability

All code is deposited on GitHub at https://gitlab.gwdg.de/tchen1/dnatensionprobe. In particular, the depository contains:

- *Matlab* (v. 2022b, MathWorks^®^ Inc.) code used for calculating the MIET calibration curves, extracting data from .ptu files, calculating the intensity and lifetime images, constructing 3D height maps, and calculating fluorescence anisotropy images.
- A ReadMe file in the depository to explain all the analysis details.

## Acknowledgments

T. Chen and J. Enderlein acknowledge financial support by the European Research Council (ERC) for financial support via project “smMIET” (grant agreement no. 884488) under the European Union’s Horizon 2020 research and innovation program. J. Enderlein acknowledges financial support by the DFG through Germany’s Excellence Strategy EXC 2067/1-390729940. D. Wang and D. Kong acknowledge financial support by the National Natural Science Foundation of China (No. 22074068). D. Wang acknowledges the scholarship from China Scholarship Council.

## Author Contributions Statement

J.E. and T.C. conceived the project. D-X.W. prepared all DNA structures and cells. D-X.W. and T.C. performed the measurements. T.C. analyzed all the data and made all figures both in the main text and supplementary information. T.C., D-X.W., and J.E wrote the manuscript. All authors revised the manuscript.

## References

[1] Discher, D.E., Janmey, P., Wang, Y.-l.: Tissue cells feel and respond to the stiffness of their substrate. Science 310(5751), 1139–1143 (2005)

[2] Qiu, Y., Brown, A.C., Myers, D.R., Sakurai, Y., Mannino, R.G., Tran, R., Ahn, B., Hardy, E.T., Kee, M.F., Kumar, S., et al.: Platelet mechanosensing of substrate stiffness during clot formation mediates adhesion, spreading, and activation. Proceedings of the National Academy of Sciences 111(40), 14430–14435 (2014)

[3] Liu, Y., Blanchfield, L., Ma, V.P.-Y., Andargachew, R., Galior, K., Liu, Z., Evavold, B., Salaita, K.: Dna-based nanoparticle tension sensors reveal that t-cell receptors transmit defined pn forces to their antigens for enhanced fidelity. Proceedings of the National Academy of Sciences 113(20), 5610–5615 (2016)

[4] Kanchanawong, P., Shtengel, G., Pasapera, A.M., Ramko, E.B., Davidson, M.W., Hess, H.F., Waterman, C.M.: Nanoscale architecture of integrin-based cell adhesions. Nature 468(7323), 580–584 (2010)

[5] Spiess, M., Hernandez-Varas, P., Oddone, A., Olofsson, H., Blom, H., Waithe, D., Lock, J.G., Lakadamyali, M., Strömblad, S.: Active and inactive β1 integrins segregate into distinct nanoclusters in focal adhesions. The Journal of cell biology 217(6), 1929 (2018)

[6] Hu, Y., Duan, Y., Salaita, K.: Dna nanotechnology for investigating mechanical signaling in the immune system. Angewandte Chemie International Edition 62(30), 202302967 (2023)

[7] Sun, X., Hao, P., Wu, N.: Dna-based mechanical sensors for cell applications. Chemistry 5(3), 1546–1559 (2023)

[8] Glazier, R., Brockman, J.M., Bartle, E., Mattheyses, A.L., Destaing, O., Salaita, K.: Dna mechanotechnology reveals that integrin receptors apply pn forces in podosomes on fluid substrates. Nature communications 10(1), 4507 (2019)

[9] Pal, K., Tu, Y., Wang, X.: Single-molecule force imaging reveals that podosome formation requires no extracellular integrin-ligand tensions or interactions. ACS nano 16(2), 2481–2493 (2022)

[10] Zhao, B., O’Brien, C., Mudiyanselage, A.P.K., Li, N., Bagheri, Y., Wu, R., Sun, Y., You, M.: Visualizing intercellular tensile forces by dna-based membrane molecular probes. Journal of the American Chemical Society 139(50), 18182–18185 (2017)

[11] Zhang, Y., Ge, C., Zhu, C., Salaita, K.: Dna-based digital tension probes reveal integrin forces during early cell adhesion. Nature communications 5(1), 5167 (2014)

[12] Zhao, B., Li, N., Xie, T., Bagheri, Y., Liang, C., Keshri, P., Sun, Y., You, M.: Quantifying tensile forces at cell–cell junctions with a dna-based fluorescent probe. Chemical Science 11(32), 8558–8566 (2020)

[13] Baig, M.M.F.A., Lai, W.-F., Akhtar, M.F., Saleem, A., Ahmed, S.A., Xia, X.-H.: Dna nanotechnology as a tool to develop molecular tension probes for bio-sensing and bio-imaging applications: An up-to-date review. Nano-Structures & Nano-Objects 23, 100523 (2020)

[14] Vicente, F.N., Chen, T., Rossier, O., Giannone, G.: Novel imaging methods and force probes for molecular mechanobiology of cytoskeleton and adhesion. Trends in Cell Biology 33(3), 204–220 (2023)

[15] Schlichthaerle, T., Lindner, C., Jungmann, R.: Super-resolved visualization of single dna-based tension sensors in cell adhesion. Nature Communications 12(1), 2510 (2021)

[16] Brockman, J.M., Su, H., Blanchard, A.T., Duan, Y., Meyer, T., Quach, M.E., Glazier, R., Bazrafshan, A., Bender, R.L., Kellner, A.V., et al.: Live-cell super-resolved paint imaging of piconewton cellular traction forces. Nature methods 17(10), 1018–1024 (2020)

[17] Chizhik, A.I., Rother, J., Gregor, I., Janshoff, A., Enderlein, J.: Metal-induced energy transfer for live cell nanoscopy. Nature Photonics 8(2), 124–127 (2014)

[18] Wang, D.-X., Liu, B., Han, G.-M., Li, Q., Kong, D.-M., Enderlein, J., Chen, T.: Metal-induced energy transfer (miet) imaging of cell surface engineering with multivalent dna nanobrushes. ACS nano 18(7), 5409–5417 (2024)

[19] Chen, T., Karedla, N., Enderlein, J.: Measuring sub-nanometer undulations at microsecond temporal resolution with metal-and graphene-induced energy transfer spectroscopy. Nature Communications 15(1), 1789 (2024)

[20] Enderlein, J.: Single-molecule fluorescence near a metal layer. Chemical Physics 247(1), 1–9 (1999)

[21] Yu, C.-h., Rafiq, N.B.M., Krishnasamy, A., Hartman, K.L., Jones, G.E., Bershadsky, A.D., Sheetz, M.P.: Integrin-matrix clusters form podosome-like adhesions in the absence of traction forces. Cell reports 5(5), 1456–1468 (2013)

[22] Chowdhury, F., Li, I.T., Leslie, B.J., Dõganay, S., Singh, R., Wang, X., Seong, J., Lee, S.-H., Park, S., Wang, N., et al.: Single molecular force across single integrins dictates cell spreading. Integrative Biology 7(10), 1265–1271 (2015)

[23] Li, H., Zhang, C., Hu, Y., Liu, P., Sun, F., Chen, W., Zhang, X., Ma, J., Wang, W., Wang, L., et al.: A reversible shearing dna probe for visualizing mechanically strong receptors in living cells. Nature Cell Biology 23(6), 642–651 (2021)

[24] Paik, D.H.: Overstretching dna at 65 pn does not require peeling from free ends or nicks. Biophysical Journal 100(3), 74 (2011)

[25] Soleimaninejad, H., Chen, M.Z., Lou, X., Smith, T.A., Hong, Y.: Measuring macromolecular crowding in cells through fluorescence anisotropy imaging with an aie fluorogen. Chemical Communications 53(19), 2874–2877 (2017)

[26] Brockman, J.M., Blanchard, A.T., Pui-Yan, V., Derricotte, W.D., Zhang, Y., Fay, M.E., Lam, W.A., Evangelista, F.A., Mattheyses, A.L., Salaita, K.: Mapping the 3d orientation of piconewton integrin traction forces. Nature methods 15(2), 115–118 (2018)

[27] Iqbal, A., Arslan, S., Okumus, B., Wilson, T.J., Giraud, G., Norman, D.G., Ha, T., Lilley, D.M.: Orientation dependence in fluorescent energy transfer between cy3 and cy5 terminally attached to double-stranded nucleic acids. Proceedings of the National Academy of Sciences 105(32), 11176–11181 (2008)

[28] Linder, S., Cervero, P., Eddy, R., Condeelis, J.: Mechanisms and roles of podosomes and invadopodia. Nature Reviews Molecular Cell Biology 24(2), 86–106 (2023)

[29] Ghosh, A., Sharma, A., Chizhik, A.I., Isbaner, S., Ruhlandt, D., Tsukanov, R., Gregor, I., Karedla, N., Enderlein, J.: Graphene-based metal-induced energy transfer for sub-nanometre optical localization. Nature Photonics 13(12), 860–865 (2019)

