## Supplementary information for "Super-Resolution Axial Imaging for Quantifying Piconewton Traction Forces in Live-cells"

**Supplementary Note 1. Determination of fluorescence lifetimes, quantum yields, and orientations of fluorophores for calculating the MIET calibration curves.**

To determine the fitting parameters for MIET calculation, the free-space lifetime $\tau_0$ was measured using the samples (Cy3-MTP, FAM-MTP, and CellMask Deep Red (CMDR)) on a glass surface (see Supplementary Figure 3). The fluorescence quantum yields (Ø) of the fluorophores were calculated using a relative method by comparing the intensity of standard fluorescence with that of the unknown samples (Cy3-MTP and FAM-MTP).[1] For this, we used fluorescein (Ø = 0.925 in 0.1 M NaOH)[2] as the references for the FAM-MTP and tetramethylrhodamine (TMR, Ø = 0.68 in methanol)[3] as reference for the Cy3-MTP.

The orientation of fluorophore was token from the literature. The 6-carboxyfluorescein (FAM) fluorophore, a fluorescein-based dye, has been reported to exhibit a random orientation within DNA structures.[4] In contrast, the cyanine-based dye Cy3 would stack against the nucleobases of the duplex perpendicular to its long axis.[4,5] Previous reports have revealed title angles of the DNA duplex from the normal to be 21° ± 2° for podosomes and 40° ± 2° for focal adhesions. Therefore, the title angles of Cy3 were 69° and 50° from vertical for those two structures, respectively, and these values were used in calculating the MIET curves of Cy3 fluorophore. For the plasma membrane (PM) staining dye CMDR, our group has already determined that it has a random orientation.[6,7] All parameters used for calculating the MIET curves are listed in Supplementary Table 2 and the calculated MIET curves are shown in Supplementary Figure 4.

**Supplementary Note 2. No FRET between Cy3-MTP and Cellmask Deep Red on the cell.**

The overlap between the fluorescence spectrum of Cy3 and absorption spectrum of CMDR would potentially induce FRET between these two dyes if the distance between them is less than 10 nm. However, in our systems, this effect is not present because the distance between Cy3 and CMDR is sufficiently larger. It has been reported that the extracellular domain of integrin adopts an upright conformation upon activation[8] and stands approximately 20 nm above the membrane surface[9]. Additionally, the DNA duplex between the dye and RGD is approximately 8.2 nm in length. Even if the DNA duplex has some tile, the substantial distance between the dye and the membrane ensures that FRET not occur in our system.

To experimentally confirm our estimation, we compared the podosome force maps using Cy3-MTPs with CMDR and without CMDR for one same cell. After collecting the Cy3 signal from one cell, we added CMDR to the observation chamber, carefully washed the chamber, and then collected the Cy3 signal at the same position. As shown in Supplementary Figure 5, the height results are nearly identical for both measurements.

It is important to note that the lifetimes and quantum yields of fluorophores are sensitive to the DNA structures,[3,5,10] we strongly recommend that anyone calculating the MIET curves measure the free-space lifetime and quantum yields specific to their own systems prior to data quantification.

**Supplementary Note 3. Lifetime determination based on the maximum likelihood estimation**

The fluorescence lifetime value for each pixel was determined by fitting the tail (starting 0.3 ns after maximum) of the TCSPC curves using a maximum likelihood estimation (MLE) method. MLE estimates decay times by minimizing the likelihood function.[11,12] It has been demonstrated that MLE is a robust method for precisely estimating fluorescence lifetime, even for decays with total counts of less than 200. Specifically, MLE provides a 20% standard error even at 20 total counts, less than 10% error at 200 total counts, and approximately 2% error when total counts > 1000.[11–13] In our measurement, the minimum counts per pixel were approximately 400 for FA measurement, ~800 for podosome measurement, and ~ 1500 for PM measurement (See supplementary Figure 9).

To evaluate the accuracy of our fitting method at different counts, we measured a standard fluorophore sample with a well-known fluorescence lifetime (atto 655 in PBS, $\tau$ = 1.80 ns).[14] We then divided the total recorded photons into bunches of different photon counts (N = 200, 400, 1000, and 7000, Supplementary Figure 8). The corresponding TCSPC curves were fitted using our MLE method, and their deviations were compared to the standard lifetime. As shown in Supplementary Figure 8, the fitted lifetime value at 400 total counts was approximately 4% lower than the expected value, with a standard error of around 8% standard error (SD/mean).

To further estimate the height error resulting from the lifetime fitting at different counts, we constructed height maps at varying photon numbers per pixel by binning frames for a sample of FAM-MTP on SLB supported by a MIET substrate. This sample was homogeneous due to fluorophore diffusion. As shown in Supplementary Figure 9, at very low counts (77 counts), the calculated mean height is 16.1 ± 2.4 nm, which is only 1 nm lower than the value calculated with 1600 counts (17.1 ± 1 nm). The mean height calculated with 320 counts is 16.5 ± 1.6 nm, demonstrating the height error in our calculation (with a minimum 400 counts used) is less than 2 nm.

**Supplementary Note 4. Calculation of the fluorophore's heights in MTP on the surface of solid MIET substrate and SLB.**

The orientations of Cy3 fluorophores in the MTP attached to integrins have been determined in previous reports.[4,15] However, their orientations on solid substrate and SLB without cells are still unknown, making it necessary to determine the fluorophores' height when the MTP is in a closed state on these substrates. To determine the fluorophore's height, we used another dye, FAM, to label the same position on the MTP. Unlike Cy3, FAM does not stack with the DNA duplex and assumes a random orientation.[4] As shown in Supplementary Figure 7, the MTPs on both solid substrate and SLB exhibit very homogeneous distributions. The mean heights of the FAM fluorophore in a 4.7 pN MTP are 17.6 ± 1.7 nm for the solid substrate and 17.1 ± 1.0 nm for the SLB.

Additionally, we did not use FAM as the main fluorophore for our cellular measurements because we found that FAM is much more prone to photobleaching and is significantly dimmer than the Cy3 tension probe, which limits imaging quality.

**Supplementary Note 5. Limitations of MIET on localizing the axial tension force of integrin**

**1) Difficult to determine the percentage of open probe**

The open percent of the DNA hairpin is a critical parameter for evaluating integrin force.[4,15–17] However, in our system, determining open percentage is challenging due to the tilt angles of DNA structures on both solid substrates and SLBs, and because MIET can only resolve the vertical position. Consequently, we cannot generate a calibration curve for calculating the open percent from either fluorescence lifetime or intensity.

2) **Ensemble averaging**

The measurement of lifetime (or height) is influenced by ensemble averaging of the fluorescence of many fluorophores within the confocal focus. The observed signal is a spatial average over the size of the excitation focus (lateral region of approximately 300 nm), resulting in an averaged outcome for both open and closed DNA hairpins. This ensemble averaging tends to underestimate the lifetime value. However, unfolding probes positioned higher emit more photons, which reduces the underestimation of lifetime. We further analyzed the signal-to-noise ratio (SNR) to evaluate this effect on height mapping for different components.

**3) Signal to Noise Ratio**

The SNR analysis for all components (FA-MTP, Podosome-MTP, and PM) is detailed in Supplementary Figure 13: (i) For the FA-MTP, the mean background count is 40 counts. In the height map analysis, only pixels with counts greater than 400 (representing regions of the FA) were included. Therefore, the SNR for the FA-MTP is much greater than 10, allowing us to ignore the underestimation effect of the folding probes for FA height mapping. (ii) For the podosome-MTP, the mean background count number is 650, and the mean count in the podosome region (count > 1000) is only 1250, resulting in an SNR of approximately 2. This low SNR leads to a larger error in the lifetime calculation from the unfolding MTP. Since half of photons in the lifetime fitting come from folding MTP s, the estimated actual lifetime of the podosome region is affected. The low SNR may be due to fluorophore diffusion, minimal height increase, and a low open percentage. The fast diffusion of Cy3-MTP compensates for photobleached probe, leading to photon accumulation in the background when scanning the same area repeatedly. (iii) For the PM, the SNR is consistently greater than 10 because only the plasma membrane is labelled by the CMDR (Supplementary Figure 13).

**4) Fluorophore orientation**

Determining the orientation of the fluorophore is crucial for calculating the MIET calibration curve. However, the presence of cyanine dyes can complicate this calculation due to their tendency to stack against the DNA duplex. While previous studies have provided insight into the average tilt angle of the fluorophore,[4,15,17] determining the orientation for focal adhesions (FAs) presents additional challenges. The tilt angle of FAs varies depending on their position within the cell: FAs exert more vertical forces at the cell center, with a median DNA tilt angle of 30°, whereas they become increasingly lateral near the cell periphery, with a median DNA tilt angle of 41°. Given that our analysis focuses solely on FAs at the cell periphery, we used a

DNA tilt angle of 40° for our MIET calculation. Importantly, this 10° difference is expected to introduce only a negligible ~2 nm height error during the conversion process.

**Supplementary Table 1:** Oligonucleotide sequences used in this work are listed above.

| Name | Sequence (5' to 3') |
|---|---|
| Ligand (ssDNA FAM) | DBCO-TTT GCT GGG CTA CGT GGC GCT CTT-FAM |
| Ligand (ssDNA CY3) | DBCO-TTT GCT GGG CTA CGT GGC GCT CTT-Cy3 |
| Anchor | CGC ATC TGT GCG GTA TTT CAC TTT-biotin |
| Hairpin-4.7 pN | GTG AAA TAC CGC ACA GAT GCG TTT GTA TAA ATG TTT TTT TCA TTT ATA CTT TAA GAG CGC CAC GTA GCC CAG C |
| Hairpin-19 pN | GTG AAA TAC CGC ACA GAT GCG CGC CGC GGG CCG GCG CGC GGT TTT CCG CGC GCC GGC CCG CGG CGA AGA GCG CCA CGT AGC CCA GC |
| cDNA-4.7 pN | AAA GTA TAA ATG AAA AAA ACA TTT ATA CAA A |
| cDNA-19 pN | CG CCG CGG GCC GGC GCG CGG AAA ACC GCG CGC CGG CCC GCG GCG |

**Supplementary Table 2** Parameters used for calculating MIET calibration curves.

| | Quantum yield, ∅ | Free-space lifetime $\tau_0$ (ns) | Orientation |
|---|---|---|---|
| FAM | 0.53 | 3.3 | Random |
| Cy3 | 0.42 | 1.8 | 69° and 50° |
| CellMask Deep Red (CMDR) | 0.3 | 1.62 | Random |

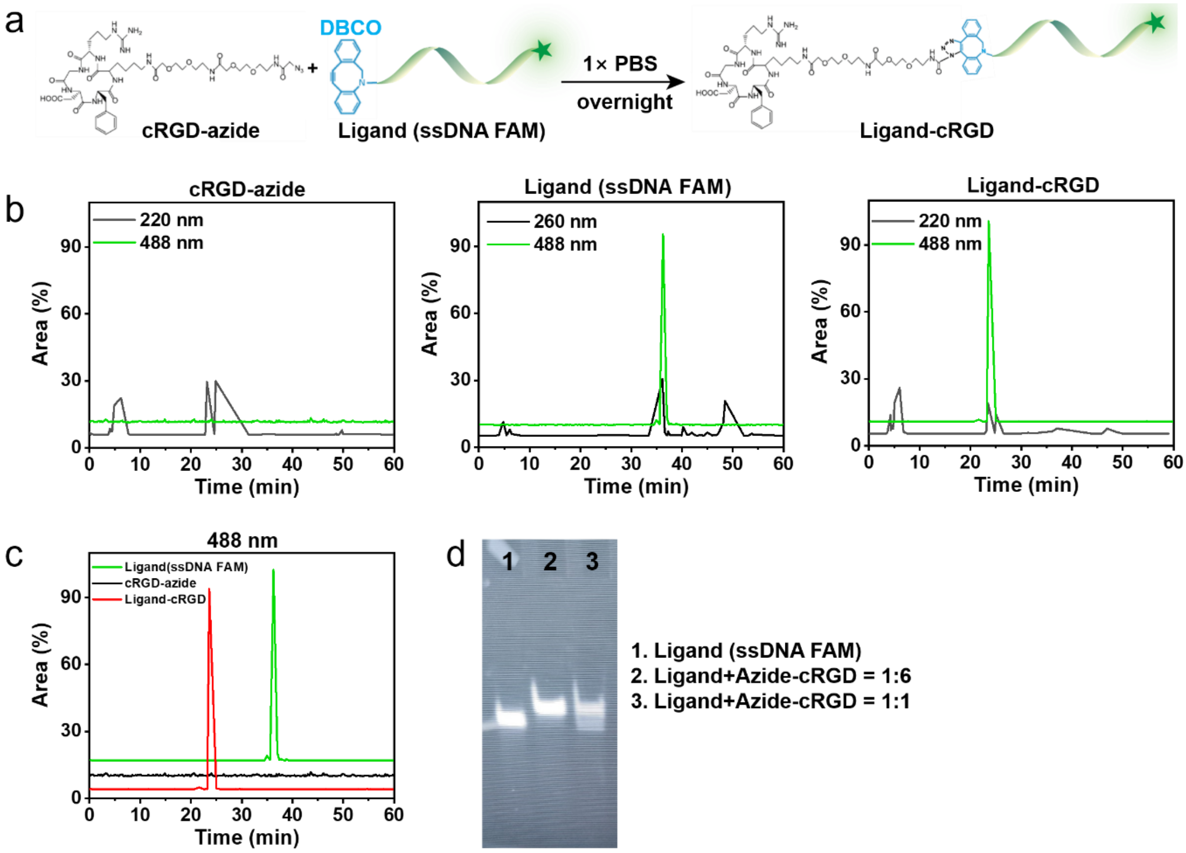


**Supplementary Figure 1. HPLC and PAGE characterization of modified probes.** (a) Schematic showing the coupling between cRGD-azide and ligand (ssDNA-FAM) to form ligand-cRGD. (b) HPLC spectra of all the starting material (cRGD and oligonucleotides) as well as the products generated in this work. (c) HPLC spectra of all oligonucleotide products under 488 nm excitation. Solvent program: 0.5 mL/min flow rate; Solvent A: 0.1 M TEAA triethylammonium acetate buffer, Solvent B: acetonitrile. Starting condition: 0-10 min 100% A; 10-40 min 0-100% gradient B. (d) 10% polyacrylamide gel electrophoresis (PAGE) characterization of modified probes. When the ratio of ligand to cRGD is 1 : 6, the reaction efficiency reaches 100%, indicating that all oligonucleotides are connected to cRGD.

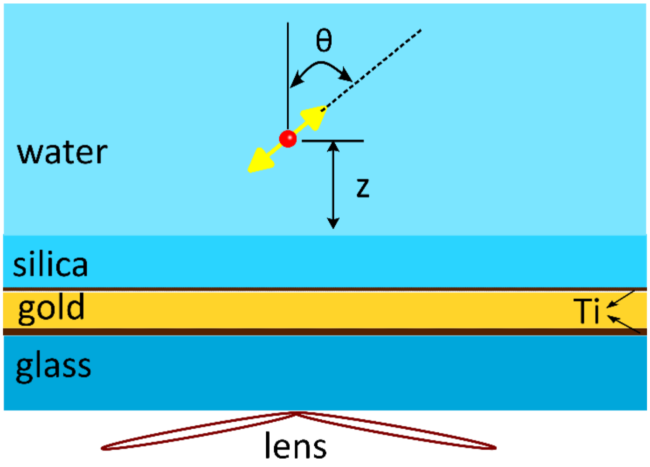


**Supplementary Figure 2. Geometry of MIET setup.** A fluorophore is positioned above a MIET substrate comprising multiple layers: 15 nm silica, 1 nm Ti, 10 nm Au, and 2 nm Ti on a commercial coverslip. Fluorescence detection and excitation are conducted using a high numerical aperture objective lens from the glass side. The fluorophore is described as an electric dipole emitter, placing at a distance z from the silica surface and its orientation is the angle θ between tis dipole axis and the optical axis.

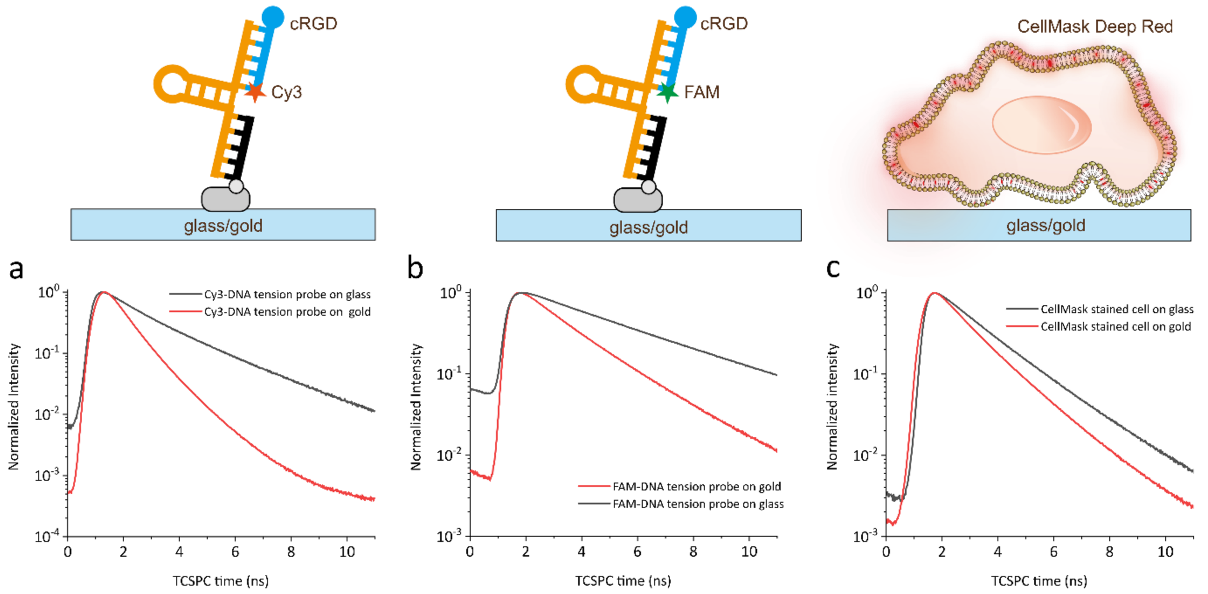


**Supplementary Figure 3. Fluorescence lifetime decay curves.** (a) Fluorescence lifetime decay curves for Cy3-MTP measured on glass surface and gold surface. (b) Fluorescence lifetime decay curves for the FAM-MTP measured on glass surface and gold surface. (c) Fluorescence lifetime decay curves for CMDR-stained cells measured on glass surface and gold surface.

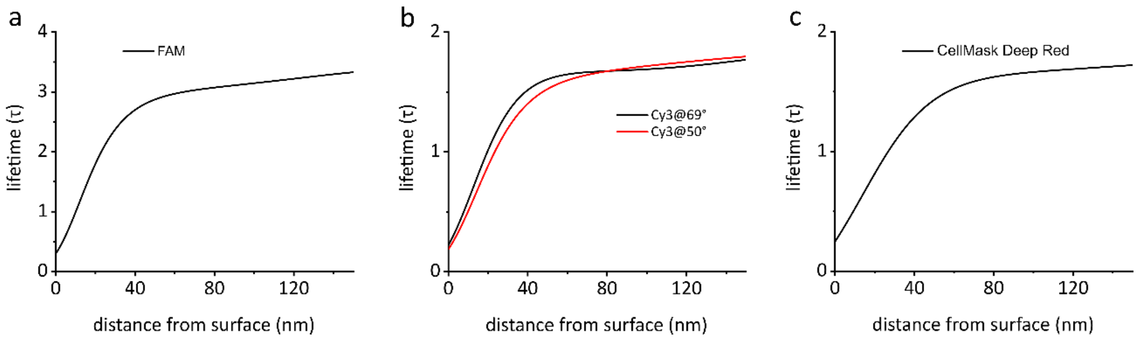


**Supplementary Figure 4. Calculated MIET calibration curves.** (a) Calculated MIET calibration curve for FAM. (B) Calculated MIET calibration curve for Cy3 with two different orientation angles. (c) Calculated MIET calibration curve for CMDR. The optical parameters of the fluorophores used for these calculations are list in Supplementary Table 2. All calculations were performed for a MIET substrate consists of 15-nm silica, 1 nm Ti, 10 nm Au, and 2 nm Ti on glass coverslip.

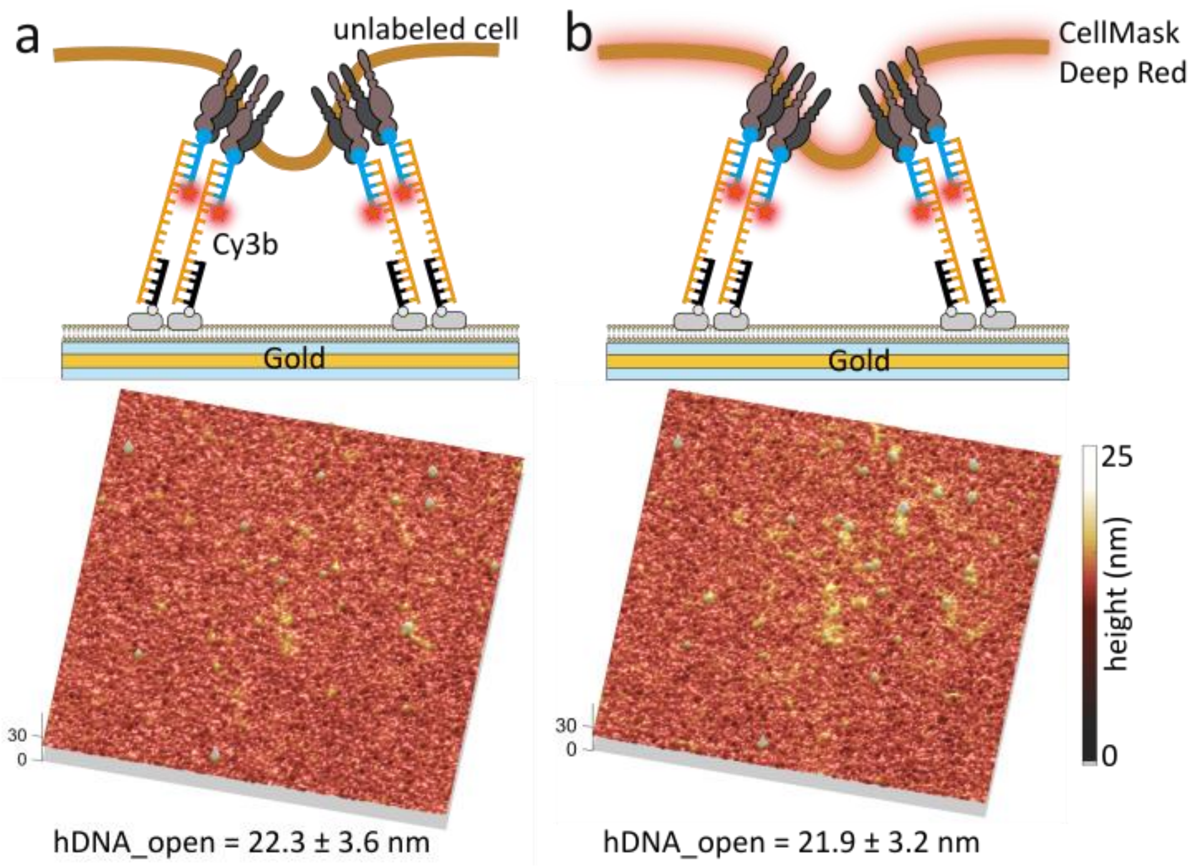


**Supplementary Figure 5. Confirmation of no FRET between Cy3-MTP and CMDR-PM.** (a) Cy3-MTP height profile for the unlabeled cell on Cy3-MTP-SLB MIET substrate. (b) Cy3-MTP height profile for the CMDR-labeled Cell on Cy3-MTP-SLB MIET substrate.

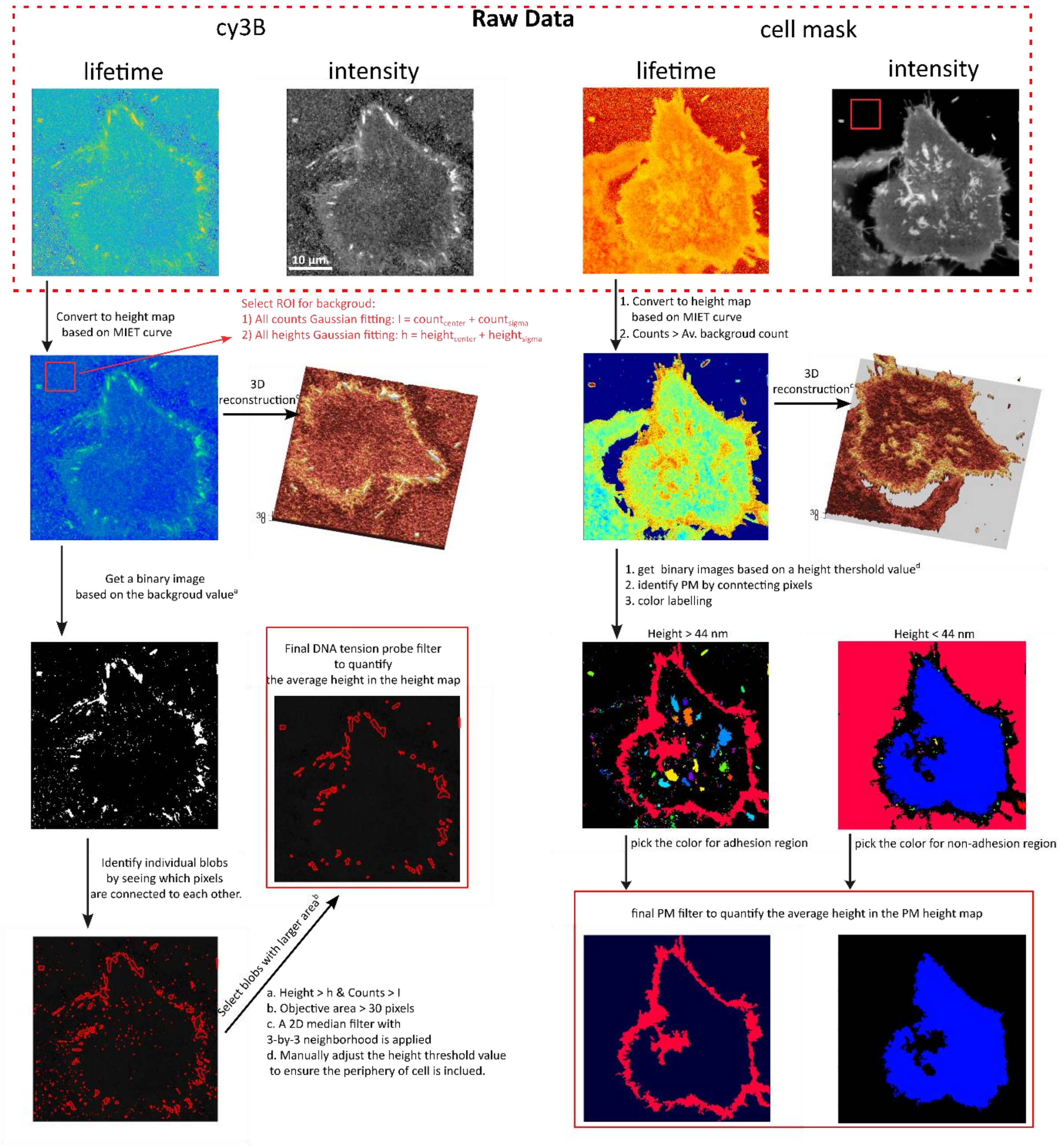


**Supplementary Figure 6: MIET analysis flowchart.** The *.ptu raw data file is processed using custom Matlab routines to generate intensity and lifetime images for each color channel. The lifetime image is then converted into height map using the lifetime-to-height calibration curve (MIET curve). To extract heights for different components (opened DNA probes, PM under adhesion, and PM under non-adhesion), segmentation and feature extraction are performed on the height maps using the Image Processing Toolbox (Image Segmentation) in Matlab. For identifying focal adhesion, the height image is first converted into a binary image based on a height threshold. Subsequently, each focal adhesion is identified by connecting the pixel that are contiguous. Blobs with small areas are excluded, resulting in an image filter specific to focal adhesions. This filter is then applied to the height map to quantify the average height of the DNA probes in the focal adhesion area. The analysis for the PM follows a similar approach, with additional steps to label connected areas with different colors and select the edge area for the adhesion region and the central area for the non-adhesion region. Finally, filters for the PM height map are obtained to quantify the average height values.

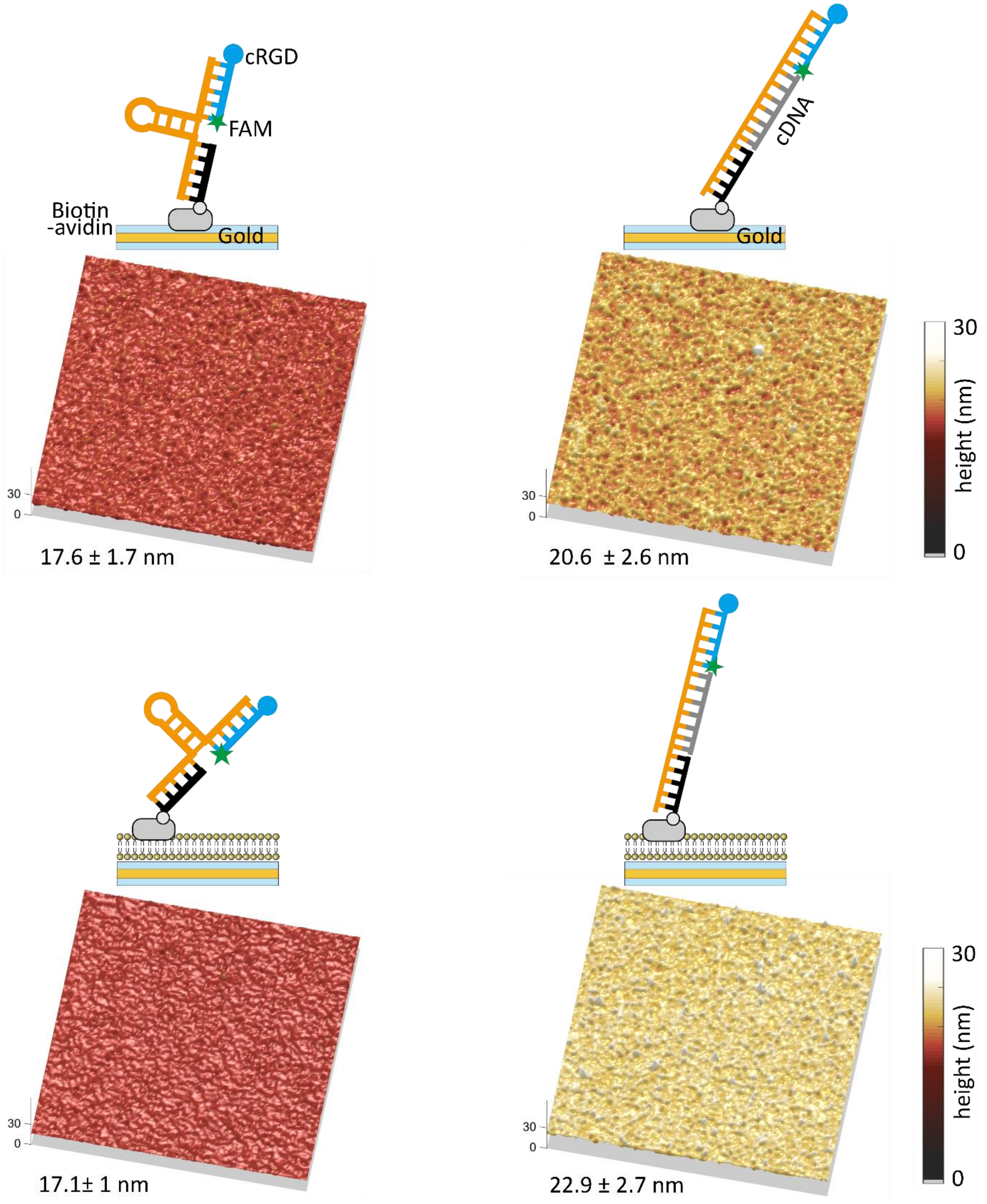


**Supplementary Figure 7. The height changes when the FAM-DNA-tension probes are unfolded with complementary strand.** Schemes and corresponding height maps showing folded (left) and unfolded (right) FAM-MTP on solid (up) MITE surface and SLB-MIET surface (bottom).

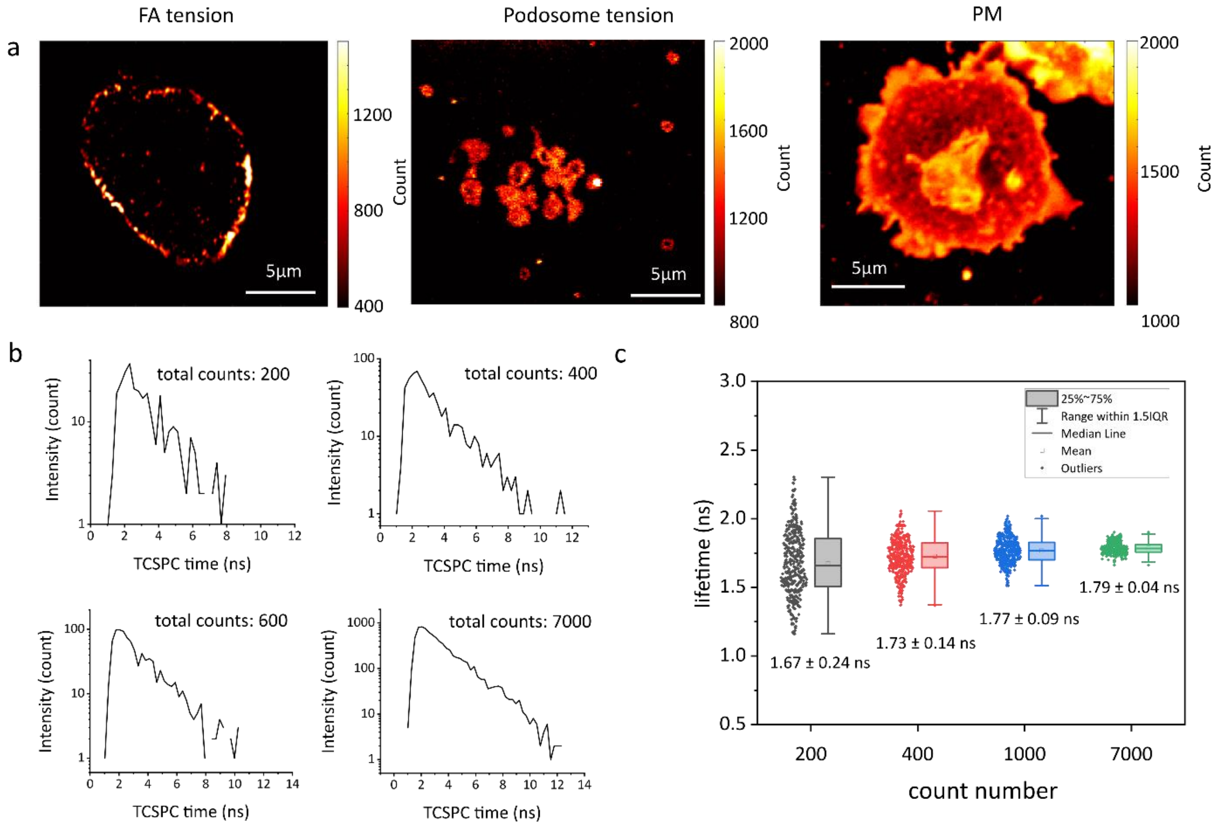


**Supplementary Figure 8: Fluorescence lifetime calculation.** (a) Representative fluorescence intensity images of FA tension, podosome tension and PM. (b) Representative TCSPC curves constructed with different photon counts for the fluorophore Atto 655 in PBS solution. (c) Fitted lifetime distributions of TCSPC curves constructed with different counts for the fluorophore Atto 655. The elements of the box plots are explained in the figure. All values are expressed as the mean ± SD. For each dataset, 400 TCSPC curves are used for fitting.

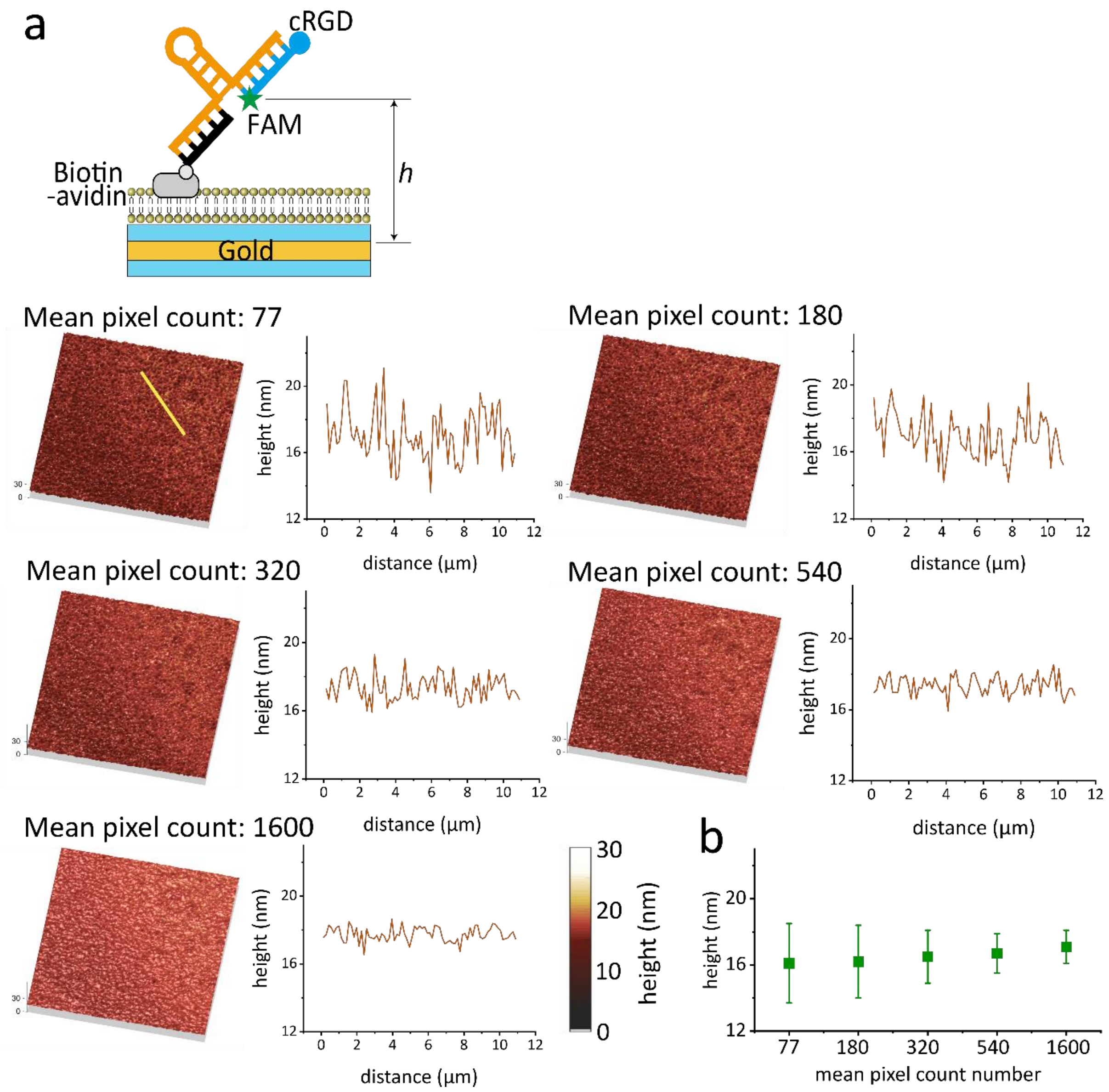


**Supplementary Figure 9: Estimation of height error based on the count numbers.** (a) The left panel displays reconstructed height maps calculated using different counts numbers, while the right panel shows the corresponding cross-section height profiles for a line marked in the first image. The color and region of color bars for all height maps are consistent. (b) The mean height values for all pixels (200 × 200) in panel (a) are depicted, with error bars representing the standard deviation.

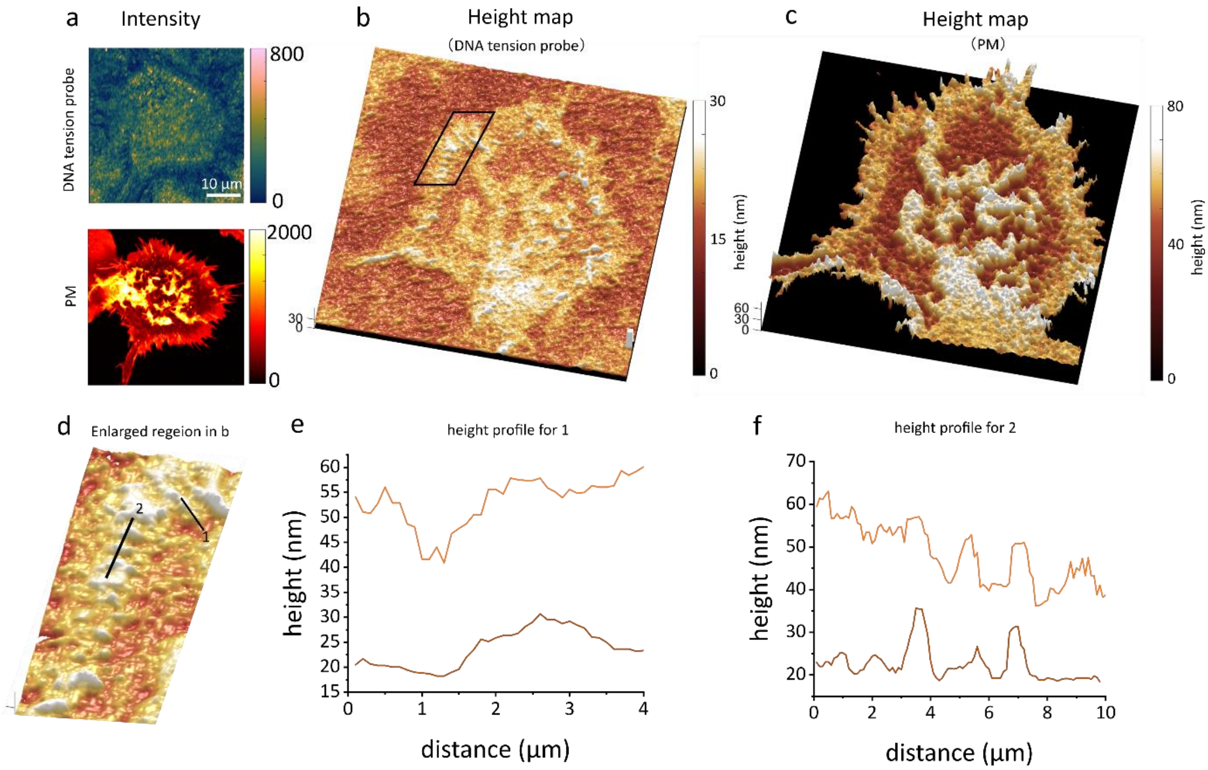


**Supplementary Figure 10. MIET measurement on 19 pN Cy3-MTP on solid substrate.** (a) Fluorescence intensity images for both Cy3-MTPs and PM. (b, c) The 3D-reconstructed height map for Cy3-MTP and PM. (d) An enlarged region marked in panel b. (e, f) Height profiles of the Cy3-MTP and PM for line 1 and line 2.

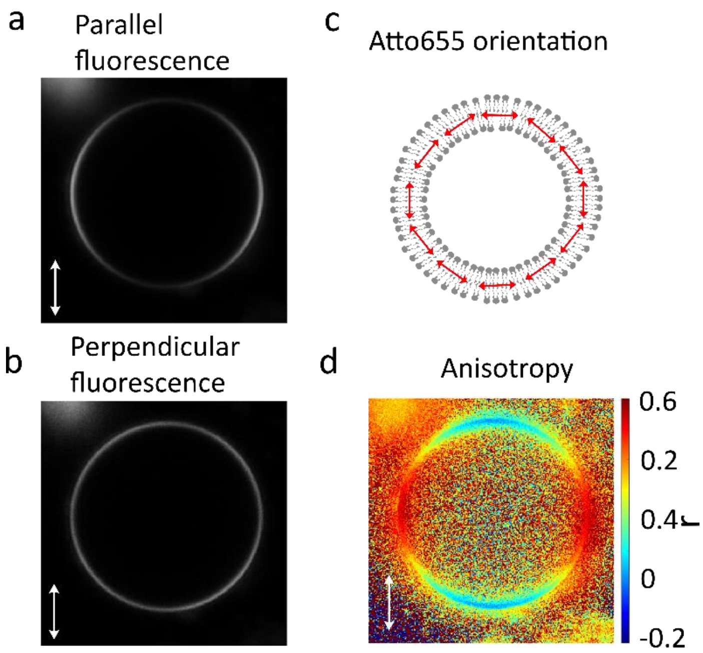


**Supplementary Figure 11: Anisotropy measurement.** (a) Representative fluorescence intensity image of Atto655-DPPE labelled GUV. The fluorescence was excited using polarized light, and the emission was collected with parallel orientation. (b) Similar as a but the emission was collected with the perpendicular orientation to the excitation polarization. (c) The orientation of Atto655-DPPE in GUV, known to align parallel to the membrane. (d) Fluorescence anisotropy image. Fluorescence anisotropy was calculated as $r = (I_{\parallel} - I_{\perp})/(I_{\parallel} + 2I_{\perp})$, where $I_{\parallel}$ and $I_{\perp}$ are the fluorescence intensities with polarization parallel and perpendicular to the excitation polarization. The results clearly demonstrate that the anisotropy image reveals the orientation of fluorophore with respect to the excitation polarization.

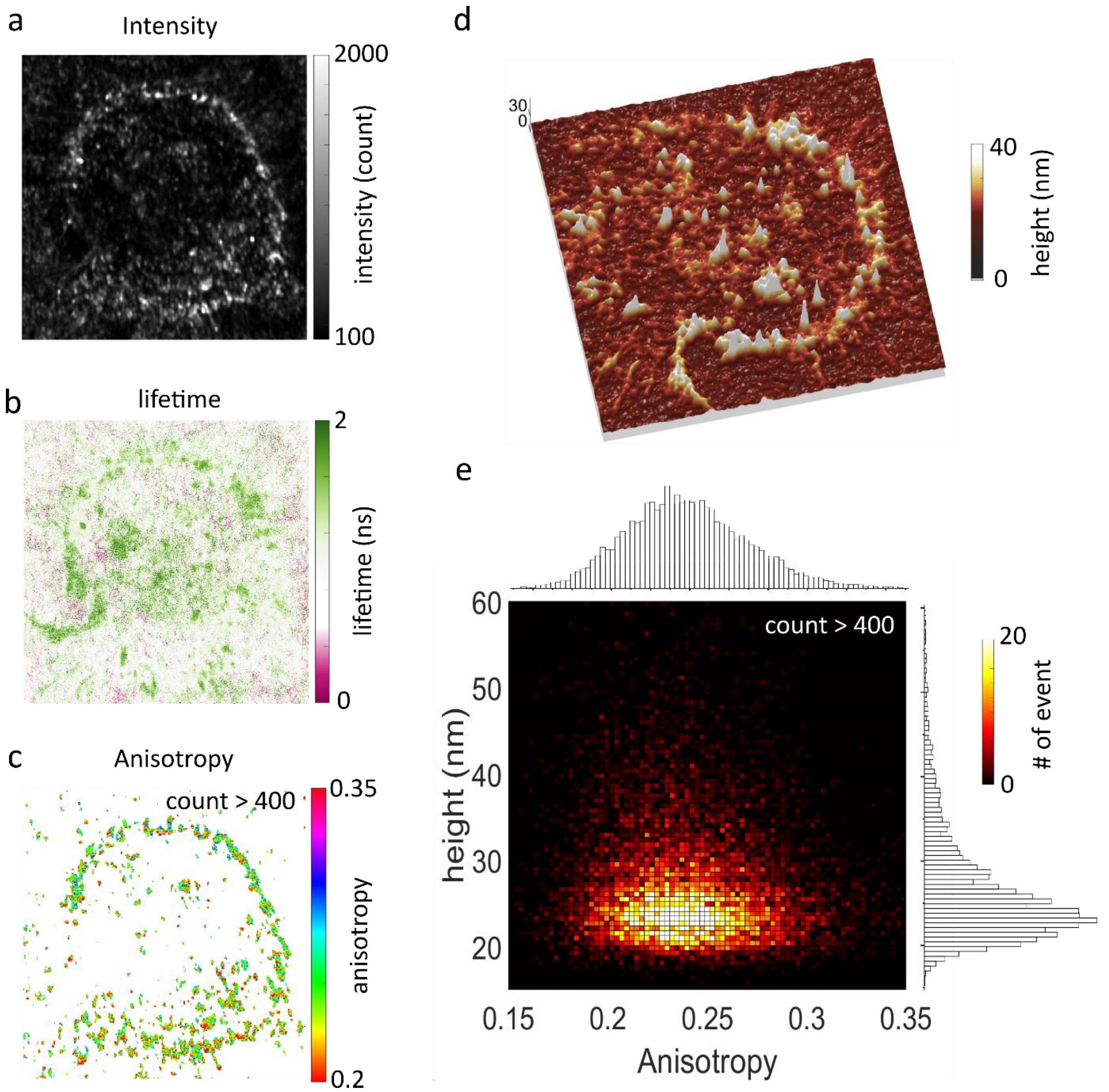


**Supplementary Figure 12. Anisotropy measurement on 19 pN Cy3-MTP on solid MIET substrate.** (a) Fluorescence intensity image of 19 pN Cy3-MTP for a Cos7 cell. (b) The corresponding lifetime image, and (c) anisotropy image. (d) 3D-reconstructed height image. (e) 2D-histograms analysis of the anisotropy versus height. The histogram is constructed by taking all the pixels values from anisotropy image and height image over the count greater than 400.

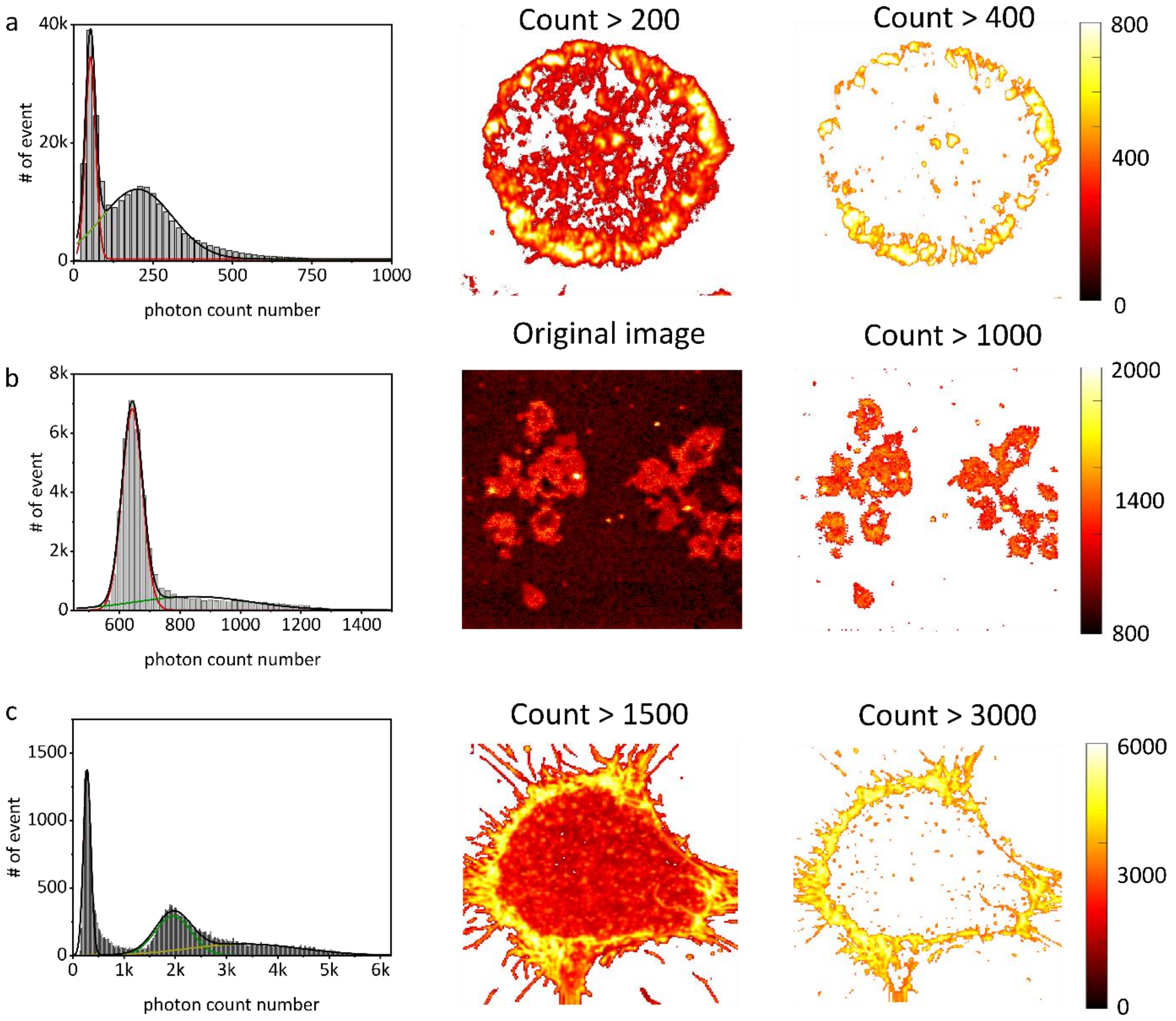


**Supplementary Figure 13. Signal-to-noise ratio analysis.** Histogram analysis of the average photon counts per pixel, along with the corresponding photon count images filtered at different count threshold for (a) FA-MTP measurement, (b) podosme-MTP measurement, and (c) PM measurement.

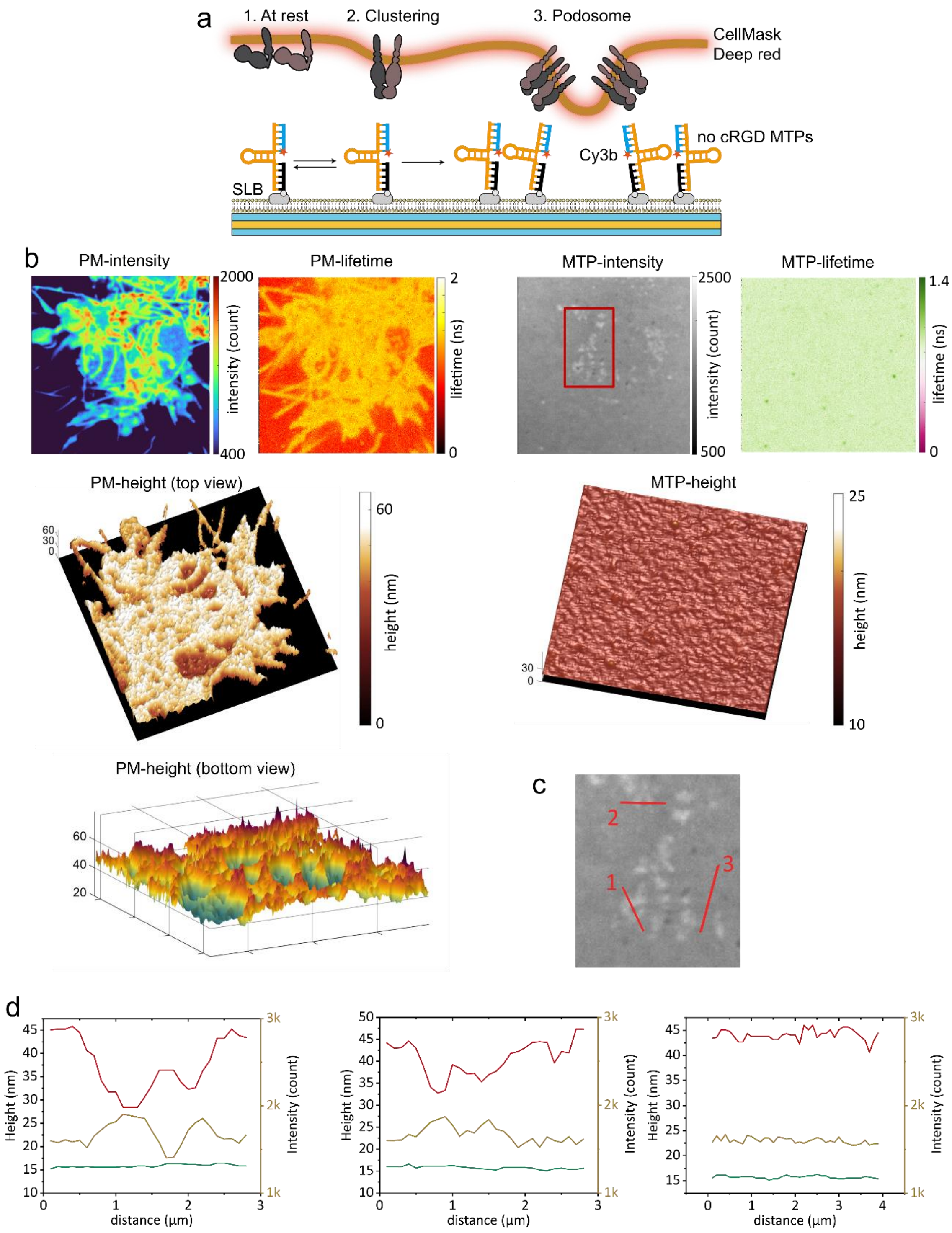


**Supplementary Figure 14. MIET measurement on 4.7 pN Cy3-MTP without cRGD on SLB.** (a) Schematic diagram of MTPs without cRGD. (b) Fluorescence intensity images, lifetime images, 3D-reconstructed height maps of the PM and 4.7 pN MTP for a single cell. (c) Enlarged area showing the marks for height profiles and intensity profiles. (d) The height profiles (red curves and green curves) for different lines marked in plane b. The yellow curves are the fluorescence intensity profile of the Cy3-DNA probe. Since the MTP was not modified with the cRGD molecule, which is necessary for integrin binding, no unfolded MTPs were observed in the lifetime or height images.

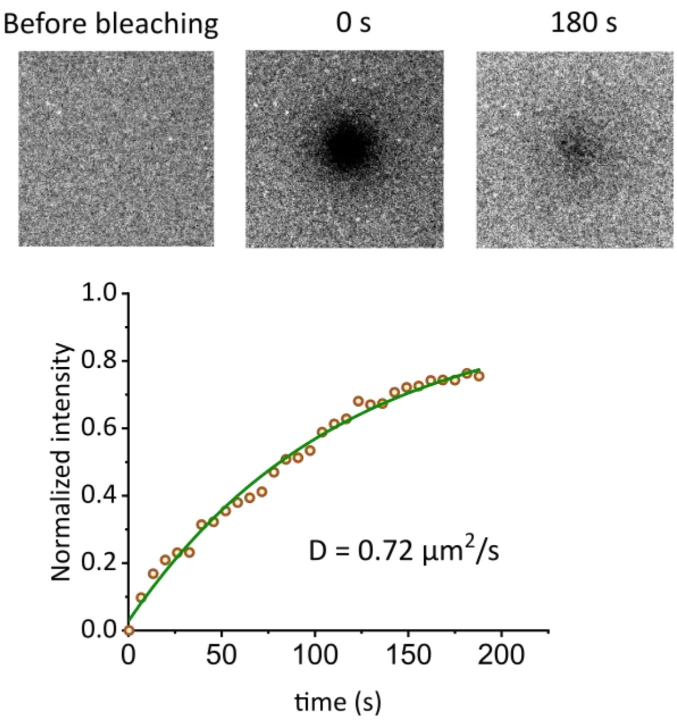


**Supplementary Figure 15: Fluorescence recovery after photobleaching (FRAP) measurement of Cy3- MTP on SLB**. TOP: representative images of Cy3-DNA-probes on SLB before photobleaching, immediately after photobleaching, and after recovery. Bottom: fluorescence recovery profile after photobleaching. Data were normalized to the SLB intensity before bleaching. Solid line was fitted with equation $y = A(1 - e^{-bx})$, where A corresponds to the mobile fraction and b is related to the diffusion time $t_{1/2} = ln2/b$. The fitting gives the diffusion coefficient of 0.72 µm$^2$/s based on the equation, $D = w^2/4t_{1/2}$, where $w$ is the radius of the bleaching area.

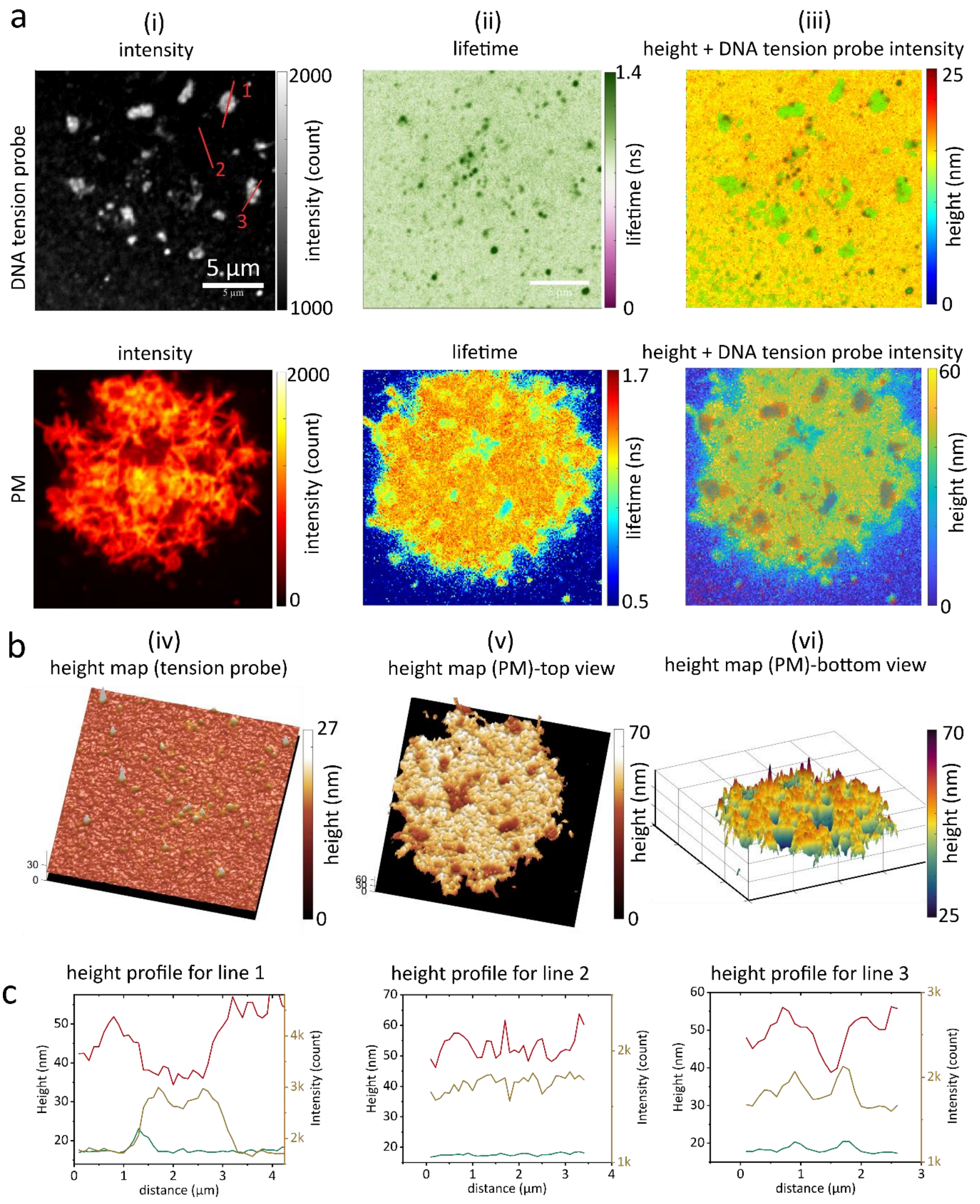


**Supplementary Figure 16 MIET measurement on 19 pN Cy3-MTP on SLB.** (a) Fluorescence intensity images (i), lifetime images (ii), and height images (iii) for Cy3-MTP and PM measured from a Cos7 cell. The height images are overlapped with the intensity image of tension probe to show that the podosome regions have higher Cy3-MTP position and lower PM height. (b) The 3D reconstructed height maps for Cy3-MTP and PM. (c) The height profiles (red curves and green curves) for different lines marked in plane a. The yellow curves are the fluorescence intensity profile of the Cy3-MTP.

**References**


1. Albrecht, C. Joseph R. Lakowicz: Principles of fluorescence spectroscopy, 3rd Edition. *Anal. Bioanal. Chem.* **390**, 1223–1224 (2008).
2. Magde, D., Wong, R. & Seybold, P. G. Fluorescence quantum yields and their relation to lifetimes of rhodamine 6G and fluorescein in nine solvents: improved absolute standards for quantum yields. *Photochem. Photobiol.* **75**, 327–334 (2002).
3. Jo Harvey, B., Perez, C. & Levitus, M. DNA sequence-dependent enhancement of Cy3 fluorescence. *Photochem. Photobiol. Sci.* **8**, 1105–1110 (2009).
4. Brockman, J. M. *et al.* Mapping the 3D orientation of piconewton integrin traction forces. *Nat. Methods* **15**, 115–118 (2018).
5. Iqbal, A. *et al.* Orientation dependence in fluorescent energy transfer between Cy3 and Cy5 terminally attached to double-stranded nucleic acids. *Proc. Natl. Acad. Sci.* **105**, 11176–11181 (2008).
6. Chizhik, A. I., Rother, J., Gregor, I., Janshoff, A. & Enderlein, J. Metal-induced energy transfer for live cell nanoscopy. *Nat. Photonics* **8**, 124–127 (2014).
7. Ghosh, A., Chizhik, A. I., Karedla, N. & Enderlein, J. Graphene- and metal-induced energy transfer for single-molecule imaging and live-cell nanoscopy with (sub)-nanometer axial resolution. *Nat. Protoc.* **16**, 3695–3715 (2021).
8. Shattil, S. J., Kim, C. & Ginsberg, M. H. The final steps of integrin activation: the end game. *Nat. Rev. Mol. Cell Biol.* **11**, 288–300 (2010).
9. Hanein, D. & Volkmann, N. Conformational Equilibrium of Human Platelet Integrin Investigated by Three-Dimensional Electron Cryo-Microscopy. *Subcell. Biochem.* **87**, 353–363 (2018).

10. Sanborn, M. E., Connolly, B. K., Gurunathan, K. & Levitus, M. Fluorescence Properties and Photophysics of the Sulfoindocyanine Cy3 Linked Covalently to DNA. *J. Phys. Chem. B* **111**, 11064–11074 (2007).
11. Maus, M. *et al.* An Experimental Comparison of the Maximum Likelihood Estimation and Nonlinear Least-Squares Fluorescence Lifetime Analysis of Single Molecules. *Anal. Chem.* **73**, 2078–2086 (2001).
12. Santra, K. *et al.* What Is the Best Method to Fit Time-Resolved Data? A Comparison of the Residual Minimization and the Maximum Likelihood Techniques As Applied to Experimental Time-Correlated, Single-Photon Counting Data. *J. Phys. Chem. B* **120**, 2484–2490 (2016).
13. Liu, X. *et al.* Fast fluorescence lifetime imaging techniques: A review on challenge and development. *J. Innov. Opt. Health Sci.* **12**, 1930003 (2019).
14. Sakhapov, D., Gregor, I., Karedla, N. & Enderlein, J. Measuring Photophysical Transition Rates with Fluorescence Correlation Spectroscopy and Antibunching. *J. Phys. Chem. Lett.* **13**, 4823–4830 (2022).
15. Glazier, R. *et al.* DNA mechanotechnology reveals that integrin receptors apply pN forces in podosomes on fluid substrates. *Nat. Commun.* **10**, 4507 (2019).
16. Zhang, Y., Ge, C., Zhu, C. & Salaita, K. DNA-based digital tension probes reveal integrin forces during early cell adhesion. *Nat. Commun.* **5**, 5167 (2014).
17. Blanchard, A. *et al.* Turn-key mapping of cell receptor force orientation and magnitude using a commercial structured illumination microscope. *Nat. Commun.* **12**, 4693 (2021).
